# Riluzole shifts glial responses to protect synapses and memory in Aβ oligomer–treated rats

**DOI:** 10.64898/2025.12.26.696626

**Authors:** Min-Kaung Wint-Mon, H Kida, R Kimura, D. Mitsushima

## Abstract

Soluble Aβ_1-42_ (amyloid beta) oligomers are potent neurotoxins that disrupt synaptic function, alter glial responses, and lead to memory impairment in Alzheimer’s disease (AD). Riluzole, a glutamate modulator approved for amyotrophic lateral sclerosis, reduces neuronal hyperexcitability, yet its *in vivo* effects on Aβ oligomer–induced cognitive dysfunction and glial alterations remain incompletely understood. Here, we investigated whether riluzole ameliorates Aβ_1-42_ oligomer–induced memory impairment and associated hippocampal pathology.

Aβ_1-42_ oligomers were bilaterally microinjected into the dorsal CA1 region of rats, followed by daily riluzole administration for seven days. Behavioral analyses revealed that riluzole significantly improved hippocampus-dependent memory, including contextual learning and spatial working memory, without affecting locomotor activity, anxiety-like behavior, or pain sensitivity. At the cellular level, riluzole reduced neuronal Aβ accumulation and attenuated synaptic and dendritic pathology, while increasing Aβ association with astrocytes and microglia in parallel with enhanced CD11b-related recognition and lysosomal processing. Notably, reductions in neuronal Aβ burden were strongly associated with improved learning performance across individual animals. Riluzole also attenuated Aβ-induced neuronal apoptosis, dendritic degeneration, and dendritic spine loss across the dorsal CA1 sublayers, and limited complement-dependent synaptic elimination by microglia.

Although Aβ oligomers induced robust recruitment of astrocytes and microglia, riluzole did not suppress glial activation; instead, it promoted morphological remodeling and shifted both astrocytes and microglia toward phenotypes associated with neuroprotection. Together, these findings indicate that coordinated remodeling of glial responses links glutamatergic modulation to enhanced amyloid handling, synaptic stability, and recovery of hippocampus-dependent memory under Aβ oligomer challenge.

**Significance Statement:** Soluble Aβ oligomers impair memory by disrupting synapses and altering glial responses in Alzheimer’s disease. Although glial activation is often viewed as detrimental, our findings demonstrate that the functional state of glia is a critical determinant of synaptic and cognitive outcomes. Using an Aβ oligomer–based rat model, we show that the glutamate modulator riluzole restores hippocampus-dependent memory not by suppressing glial recruitment, but by promoting coordinated functional and morphological remodeling of astrocytes and microglia toward phenotypes associated with neuroprotection. These changes are accompanied by enhanced glia-mediated amyloid handling, reduced complement-dependent synaptic elimination, and preservation of dendritic spine integrity. Across individual animals, hippocampus-dependent learning performance was negatively associated with neuronal Aβ burden, linking glial-mediated amyloid handling to behavioral recovery.

## Introduction

The aggregation and accumulation of the β-amyloid (Aβ) peptide are central features of Alzheimer’s disease (AD) pathogenesis (Hardy & Selkoe, 2002). While early formulations of the amyloid hypothesis emphasized the role of insoluble plaques, subsequent work identified soluble Aβ_1-42_ oligomers as key neurotoxic species that initiate synaptic dysfunction and cognitive decline (Hardy & Selkoe, 2002; Hector & Brouillette, 2021). Consistent with this view, intracerebral administration of natural or synthetic Aβ_1-42_ oligomers induces synaptic loss, astrogliosis, neuronal death, and memory impairment in rodents (Brouillette et al., 2012; Forny-Germano et al., 2014). These findings position soluble Aβ_1-42_ oligomers as a principal pathogenic driver of AD-related cognitive dysfunction.

A growing body of evidence indicates that soluble Aβ oligomers profoundly disrupt glutamatergic signaling, leading to neuronal hyperexcitability and network instability (Dickerson et al., 2005; Tu et al., 2014; Zott et al., 2019). Such glutamatergic dysregulation impairs long-term potentiation (LTP) and disturbs the excitatory/inhibitory (E/I) balance, ultimately compromising learning and memory (Lei et al., 2016; Yang et al., 2021). In parallel, excessive extracellular glutamate and overactivation of glutamatergic receptors promote Ca²⁺ overload, synaptic damage, and neuronal death (Li et al., 2011; Hascup & Hascup, 2016; Calvo-Rodriguez et al., 2020). Together, these observations underscore glutamatergic dysfunction as a critical and therapeutically relevant mechanism in Aβ-induced pathology.

Riluzole (2-amino-6-(trifluoromethoxy)benzothiazole), an FDA-approved treatment for amyotrophic lateral sclerosis (ALS), is a well-characterized glutamate modulator that reduces presynaptic glutamate release and enhances glutamate uptake (Fumagalli et al., 2008; Hascup et al., 2021). Through these actions, riluzole stabilizes excitatory network activity and improves synaptic plasticity, including LTP (Yang et al., 2021; Shen et al., 2025). In addition to its effects on neuronal excitability, riluzole has been shown to reverse age-associated transcriptional changes, synaptic alterations, and memory decline (Pereira et al., 2014, 2017), highlighting its broad neuroprotective potential (Zarate & Manji, 2008; Lamanauskas & Nistri, 2008). These properties suggest that riluzole may be well suited to counteract Aβ-driven circuit dysfunction in AD.

Neuronal hyperexcitability is closely linked to cognitive impairment in AD (Dickerson et al., 2005; Koh et al., 2010) and has been attributed in part to alterations in the persistent sodium current (I_NaP) (Ren et al., 2014). We previously demonstrated that riluzole suppresses Aβ_1-42_ oligomer–induced neuronal hyperexcitability in vitro by inhibiting I_NaP (Min-Kaung-Wint-Mon et al., 2024). Moreover, riluzole improves cognitive performance and reduces amyloid burden in transgenic models of AD (Hunsberger et al., 2015; Okamoto et al., 2018; Hascup et al., 2021). However, whether riluzole can ameliorate Aβ oligomer–induced cognitive dysfunction *in vivo*, and how it affects the cellular mechanisms linking glutamatergic dysregulation to synaptic and circuit pathology, remain incompletely understood.

In the present study, we investigated the effects of riluzole in a rat model of Aβ_1-42_ oligomer–induced hippocampal dysfunction. We tested the hypothesis that riluzole rescues memory impairment not simply by dampening neuronal hyperexcitability, but by reshaping glial responses to Aβ pathology. Specifically, we examined whether riluzole preserves synaptic integrity and cognitive function by promoting neuroprotective functional states of astrocytes and microglia that support amyloid handling and limit maladaptive synaptic remodeling.

## Materials and Methods

### Animals

Male Sprague–Dawley rats (4 weeks old; Chiyoda Kaihatsu, Japan) were housed under standard laboratory conditions (23 ± 1°C, 12 h light/dark cycle) with ad libitum access to food and water. Only male Sprague–Dawley rats were used in this study to reduce variability associated with estrous cycle–dependent hormonal fluctuations. Estrogen has been reported to modulate Aβ-induced neurotoxicity and cognitive impairment (Kim et al., 2022); therefore, restricting the experimental cohort to males allowed more precise assessment of the effects of Aβ_1-42_ oligomers and riluzole. The potential influence of sex differences is acknowledged as a limitation and warrants future investigation.

A total of 35 rats were used, and all procedures were conducted in accordance with institutional guidelines to minimize animal suffering.

### Preparation of Aβ_1-42_ oligomers

Synthetic Aβ_1-42_ peptide was prepared according to a previously established oligomerization protocol (Stine et al., 2003), with minor modifications. Briefly, lyophilized Aβ_1-42_ peptide was dissolved in hexafluoro-2-propanol (HFIP) to disaggregate pre-existing assemblies, dried under vacuum, and reconstituted in dimethyl sulfoxide (DMSO). The peptide solution was then diluted in phosphate-buffered saline (PBS) to obtain an oligomer-enriched preparation and incubated at 4 °C for 24 h. Aliquots were stored at −40 °C until use and diluted in 0.9% NaCl immediately before microinjection. The final DMSO concentration was 0.4%. The oligomeric composition of the preparation was verified by SDS–PAGE and immunoblotting using lecanemab and 6E10 antibodies (Fig. 1B).

**Figure 1.**
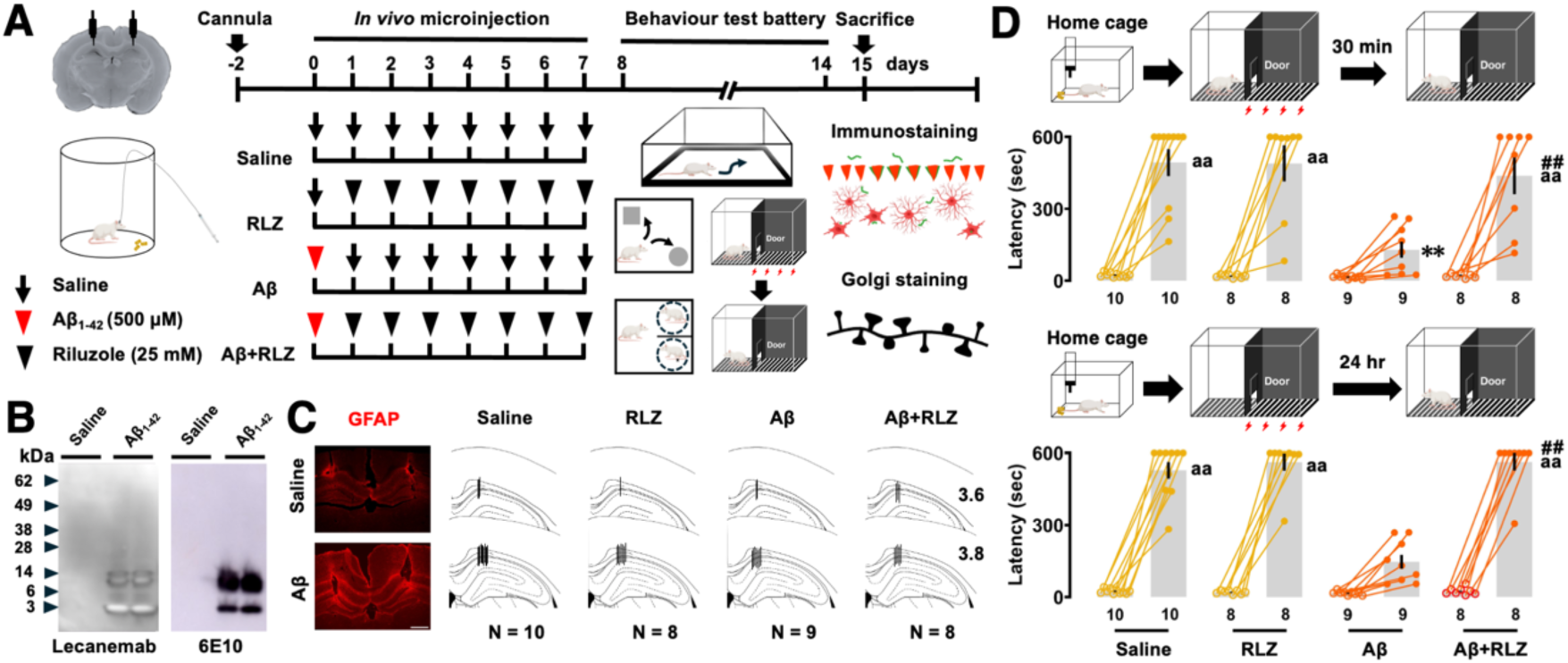
Riluzole attenuates Aβ_1-42_ oligomer–induced contextual memory impairment in the inhibitory avoidance (IA) task. (A) Experimental timeline and schematic of *in vivo* microinjection. A guide cannula was implanted above the dorsal CA1 region, and rats received bilateral microinjection of Aβ_1-42_ oligomers, followed by daily riluzole administration for 7 days. Behavioral testing was conducted before tissue collection for histological analyses. (B) Representative immunoblots of the Aβ_1-42_ oligomer preparation separated by 4–20% gradient SDS-PAGE and probed with anti-Aβ antibodies (lecanemab and 6E10). (C) Representative GFAP immunofluorescence images showing astrocytic responses after Aβ_1-42_ oligomer injection (left) and schematic reconstructions of microinjection sites in the dorsal hippocampus (right). Numbers indicate the distance (mm) posterior to bregma. Scale bar, 1000 µm. (D) Contextual learning assessed by the IA task. Schematic illustrations of the task are shown above the plots. Rats received a foot shock (1.6 mA, 2 s) upon entry into the dark compartment during training. Latency to enter the dark compartment was measured 30 min (top) and 24 hr (bottom) after training. Open circles indicate training trials and closed circles indicate test trials. Data are shown as individual values with mean ± SEM; sample sizes are indicated below each group. ^aa^*P* < 0.01 *vs* training; \*\**P* < 0.01 *vs* saline; ^##^*P* < 0.01 *vs* Aβ. Illustrations of rats were created using BioRender.

This preparation is referred to as ‘Aβ oligomers’ throughout the manuscript, and all experiments were performed using the same preparation batch.

### Western blotting

Samples were prepared by mixing Aβ_1-42_ peptide (100 µM) with 2× Laemmli sample buffer at a 1:1 ratio. Aliquots (10 µl) were separated on 4–20% gradient SDS–PAGE gels (Mini-PROTEAN TGX, Bio-Rad) and transferred onto PVDF membranes (0.45 µm). Membranes were blocked with 5% skim milk in TBS-T and incubated overnight at 4 °C with lecanemab (1:5000; A3112, Selleck Biotechnology) or 6E10 (1:5000; 803004, BioLegend). After washing, membranes were incubated with HRP-conjugated anti-human or anti-mouse secondary antibodies (1:5000) for 1 h at room temperature. Immunoreactive bands were visualized using enhanced chemiluminescence and imaged with an Amersham Imager 680.

### Stereotaxic surgery and microinjection

Rats were anesthetized with ketamine (100 mg/kg) and xylazine (10 mg/kg) and placed in a stereotaxic frame. Stainless-steel guide cannula was implanted bilaterally above the dorsal hippocampal CA1 region (AP −3.0 mm, ML ±2.0 mm, DV −1.0 mm from dura). On the experimental day, the stylet was replaced with a 1-mm longer injector (outer diameter = 0.31 mm) through a fine, flexible silicone tubing (outer diameter = 0.5 mm, Kaneka Medix Co. Osaka, Japan). Aβ_1-42_ oligomers (500 µM, 1 µL per side) were bilaterally injected at a rate of 1 µL/5 min without restraining the animals. Riluzole (25 mM, A2423; TCI, Tokyo, Japan) was administered daily for the subsequent 7 days (**Figure 1A**). Injectors were left in place for 5 min after infusion to prevent reflux. Cannula placement was verified histologically (**Figure 1C**).

### Behavioral test battery

After riluzole treatment, rats underwent a behavioral test battery to assess hippocampus-dependent memory and basic physiological functions. Animals were habituated to the testing environment, and behavior was recorded by video. Apparatuses were cleaned between trials, and a 30-min inter-test interval was maintained (Sakimoto et al., 2022; Min-Kaung-Wint-Mon et al., 2024).

### Open field test

Locomotor activity and anxiety-related behavior were assessed by recording total distance traveled, center time, center entries, and rearing during a 5-min session.

### Object-based memory tasks

Exploration was defined as directing the nose toward an object within 2 cm or touching it with the forelimbs for ≥1 s; sitting or leaning against objects was excluded.

- Object recognition: One familiar object was replaced with a novel object during the test phase.
- Object location: One familiar object was moved to a new location.
- Object-in-place: Two of four objects were exchanged.
- Object recency: One object from the first sampling phase and one from the second were presented.

Memory performance was quantified as the ratio of exploration time for the novel object or location to total exploration time (Cruz-Sanchez et al., 2020; Sakimoto et al., 2022). We measured the time spent interacting with the objects after 5-min, 30-min and 24-hr intervals.

### Social behavior and memory

Social interaction time was measured during free interaction with an unfamiliar conspecific. Social preference and recognition were assessed using a two-choice task, and performance was expressed as the ratio of interaction time with the social or novel target to total interaction time.

### Light/dark test

Anxiety-like behavior was evaluated by measuring time spent in the lit compartment and rearing frequency (Min-Kaung-Wint-Mon et al., 2024).

### Y-maze task

Working memory was assessed by spontaneous alternation behavior during a 5-min session. After 24 hr, the rats were placed in one of the arms again, and their behavior was recorded for 5 min. Alternation percentage was calculated as (actual alternations / possible alternations) × 100 (Min-Kaung-Wint-Mon et al., 2024).

### Inhibitory avoidance (IA) task

Contextual learning was assessed by measuring latency to enter the shock-associated compartment during test sessions conducted 30 min and 24 h after training (Mitsushima et al., 2011; Min-Kaung-Wint-Mon et al., 2024).

### Fear conditioning test

Rats received three foot shocks (0.8 mA, 2 s) during training. Freezing behavior was assessed 1 h and 24 h later (Min-Kaung-Wint-Mon et al., 2024). Freezing was defined as the cessation of all behaviors except respiration for at least 1 sec.

### Flinch-jump test

Pain sensitivity was evaluated by determining flinch, jump, and vocalization thresholds in response to increasing foot shock intensities (Min-Kaung-Wint-Mon et al., 2024).

### Immunohistochemistry

Immunohistochemistry was performed as previously described (Min-Kaung-Wint-Mon et al., 2024). Briefly, brains were perfused, fixed, cryoprotected, and sectioned (20 µm). Sections were incubated with primary and secondary antibodies (**Table S1**), mounted, and imaged under identical conditions.

### Image acquisition and analysis

Images were acquired under identical conditions using fluorescence microscopy. Cell density was calculated as cells/mm², and fluorescence intensity was quantified within defined ROIs using ImageJ. Two to three sections per animal (number of rats = 4-5) were analyzed by an experimenter blind to treatment.

### Analysis of internalized Aβ and synapses

Z-stack images were acquired and reconstructed for 3D analysis using an all-in-one fluorescence microscope (BZ-X810, KEYENCE Co., Osaka, Japan). Internalized Aβ and synaptic markers within microglia were quantified using ImageJ. Synaptic colocalization was assessed by measuring vGlut1 and PSD95 puncta density within defined ROIs.

### Morphological analysis of astrocytes and microglia

Cell morphology was reconstructed from z-stack images. Soma size, territory area, process length, branching, and Sholl intersections were quantified using Fiji/ImageJ.

### Classification of microglia

Microglia were classified as ramified, hypertrophic, dystrophic, amoeboid, or rod-shaped based on established morphological criteria (Martini et al., 2020).

### Golgi-Cox staining and spine analysis

Golgi-Cox staining was performed as previously described (Zhong et al., 2019). Spine density and morphology were analyzed along 20-µm dendritic segments across the dorsal CA1 sublayers. Spines were classified as thin, mushroom, or stubby based on head and neck dimensions (Mehder et al., 2020).

### Statistical analysis

Data are presented as mean ± SEM. Normality was assessed using the Shapiro–Wilk test. Behavioral and cellular data were analyzed using two-way ANOVA with Aβ and riluzole as factors, followed by Bonferroni post hoc tests. Recognition memory tasks were analyzed using paired t-tests comparing novel versus familiar targets. The association between contextual learning and neuronal Aβ burden was assessed by Pearson’s correlation analysis. Statistical significance was set at p < 0.05.

## Results

### Effect of riluzole on Aβ_1-42_ oligomer–induced contextual memory impairment

To examine the effects of riluzole on Aβ₁–₄₂ oligomer–induced memory impairment, Aβ oligomers were bilaterally microinjected into the dorsal CA1 region, followed by daily riluzole administration for 7 days. After completion of the behavioral test battery, brains were collected for immunohistochemical and Golgi analyses (**Figure 1A**).

To characterize the Aβ_1-42_ oligomer preparation used in this study, peptides were analyzed by 4–20% gradient SDS-PAGE under non-denaturing and non-reducing conditions. Immunoblotting with lecanemab detected a mixture of low–molecular weight species (monomers to tetramers) and higher-order oligomers, whereas immunodetection with 6E10 predominantly revealed low–molecular weight Aβ species (**Figure 1B**). In addition, GFAP immunoreactivity was increased in the dorsal CA1 region following Aβ_1-42_ oligomer injection, indicating astrocytic recruitment to the injection site (**Figure 1C**).

Contextual learning was assessed using the inhibitory avoidance (IA) task (**Figure 1D**). Thirty minutes after training, a three-way ANOVA revealed significant main effects of training (*F*₁,₆₂ = 147.503, *P* < 0.0001), Aβ (*F*_1,62_ = 12.076, *P* = 0.0009), and riluzole (*F*_1,62_ = 6.675, *P* = 0.0121), as well as a significant interaction among these factors (*F*_1,62_ = 6.220, *P* = 0.0153). Post hoc analyses showed that training increased step-through latency in the saline, RLZ, and Aβ + RLZ groups, whereas this increase was not observed in the Aβ group.

When retention was reassessed 24 h after training, a three-way ANOVA again revealed significant main effects of training (*F*_1,62_ = 626.622, *P* < 0.0001), Aβ (*F*_1,62_ = 31.983, *P* < 0.0001), and riluzole (*F*_1,62_ = 44.963, *P* < 0.0001), together with a significant interaction (*F*_1,62_ = 29.094, *P* < 0.0001). Although step-through latency increased across all groups at 24 h, Aβ-treated rats exhibited reduced latency relative to controls, whereas riluzole treatment increased latency in Aβ-injected rats. Together, these results indicate that riluzole attenuates Aβ_1-42_ oligomer–induced deficits in contextual memory.

### Effect of riluzole on sensory, motor, and emotional functions

Because performance in hippocampus-dependent memory tasks can be influenced by basic physiological and emotional factors, we assessed locomotor activity, anxiety-like behavior, pain sensitivity, object exploration, and social behavior. In the open-field test, there were no significant group differences in total distance traveled (Aβ: *F*_1,31_ = 0.011, *P* = 0.9160), time spent in the center (Aβ: *F*_1,31_ = 2.459, *P* = 0.1270), latency to enter the center (Aβ: *F*_1,31_ = 0.250, *P* = 0.6206), frequency of center entries (Aβ: *F*_1,31_ = 0.204, *P* = 0.6547), or rearing activity (Aβ: *F*_1,31_ = 1.198, *P* = 0.2822), indicating that neither locomotor activity nor exploratory behavior was altered across groups (**Figure 2A**). To determine whether motivation influenced recognition performance, object exploration and social behaviors were examined. Total object exploration time did not differ among groups (**Figure 2B**; Aβ: *F*_1,31_ = 0.210, *P* = 0.6501). Similarly, no group differences were observed in total social interaction time (Aβ: *F*_1,31_ = 0.061, *P* = 0.8072) or social preference (Aβ: *F*_1,31_ = 0.661, *P* = 0.4225), indicating comparable motivation to explore objects and engage in social interactions (**Figure 2C**).

**Figure 2.**
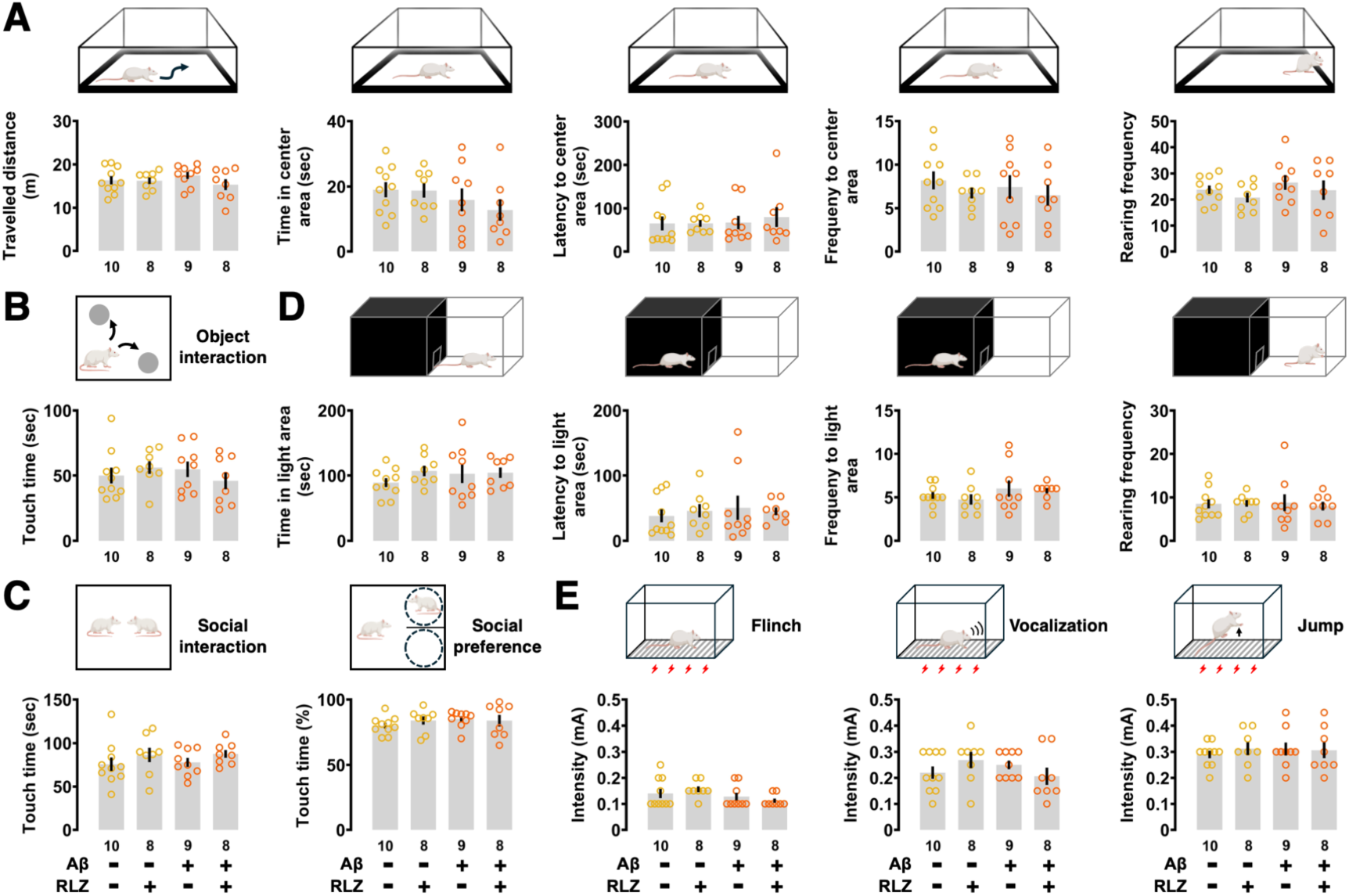
Riluzole does not alter basic locomotor, emotional, social, or sensory functions. (A) Open-field test showing comparable total distance traveled, time spent in the center, latency to enter the center, center-entry frequency, and rearing activity across groups. (B) Total object exploration time was not affected by Aβ_1-42_ oligomers or riluzole treatment. (C) Social interaction time and social preference were similar among all groups. (D) Light/dark test showing no group differences in time spent in the lit compartment, latency to enter the light area, visit frequency, or rearing activity. (E) Flinch–jump test showing unaltered current thresholds for flinch, vocalization, and jump responses. Data are shown as individual values with mean ± SEM; sample sizes are indicated below each group.

Consistent with these findings, performance in the light/dark test revealed no significant differences in time spent in the light compartment (Aβ: *F*_1,31_ = 0.305, *P* = 0.5846), latency to enter the light compartment (Aβ: *F*_1,31_ = 0.216, *P* = 0.6457), frequency of light compartment entries (Aβ: *F*_1,31_ = 2.165, *P* = 0.1512), or rearing activity (Aβ: *F*_1,31_ = 0.018, *P* = 0.8939), indicating no detectable changes in anxiety-like behavior (**Figure 2D**). Finally, pain sensitivity assessed by the flinch–jump test was not altered, as there were no significant differences in flinch (Aβ: *F*_1,31_ = 3.795, *P* = 0.0605), vocalization (Aβ: *F*_1,31_ = 0.378, *P* = 0.5432), or jump thresholds (Aβ: *F*_1,31_ = 0.100, *P* = 0.7542) (**Figure 2E**).

Together, these results indicate that riluzole does not affect sensory, motor, or emotional functions, supporting the conclusion that the observed improvements in hippocampus-dependent memory are not attributable to nonspecific behavioral confounds.

### Effect of riluzole on Aβ_1-42_ oligomer–induced impairment in hippocampus-dependent memory tasks

We next examined whether riluzole rescues deficits in multiple hippocampus-dependent memory domains using a battery of recognition, associative, social, and contextual tasks. In the object recognition task (**Figure 3A**), touch time of the novel object increased significantly in the saline (*t* = 7.288, *P* < 0.0001, paired t-test), RLZ (*t* = 6.858, *P* = 0.0002), and Aβ+RLZ groups (*t* = 4.263, *P* = 0.004). However, the Aβ group did not show an increase in touch time after a 5-min test interval (*t* = 0.155, *P* = 0.881). Aβ-induced impairment in object recognition memory improved by RLZ injection after 5 min. Thirty minutes later, the saline (*t* = 6.497, *P* = 0.0001), RLZ (*t* = 4.469, *P* = 0.0029), and Aβ+RLZ groups (*t* = 3.327, *P* = 0.013) continued to explore novel objects, while the Aβ group showed a reduced preference (*t* = 3.083, *P* = 0.015, paired t-test). Aβ-induced impairment improved with RLZ after 30 min. However, after 24 hours, all groups spent a comparable amount of time exploring both objects.

**Figure 3.**
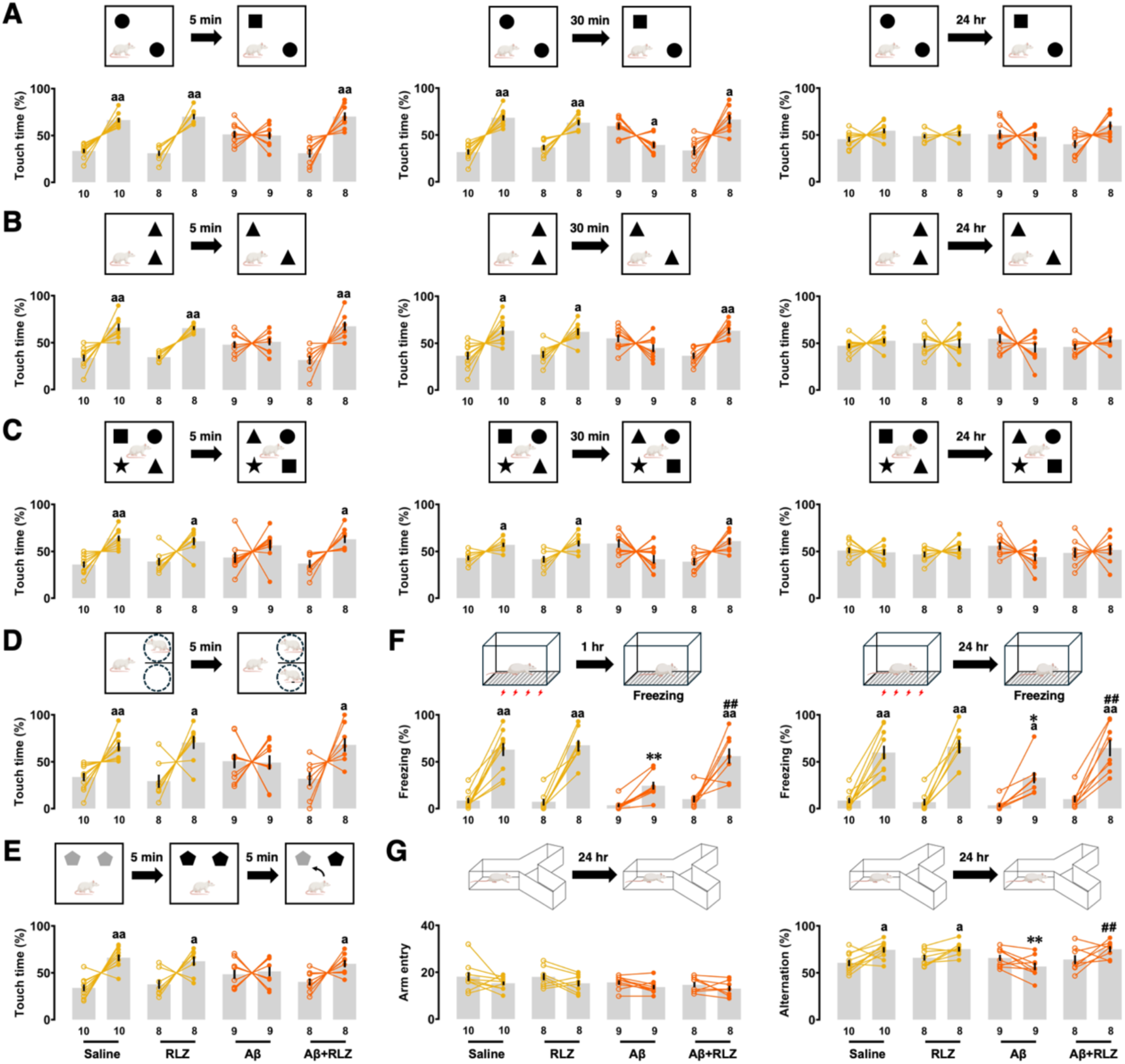
Riluzole improves Aβ_1-42_ oligomer–induced impairments across multiple hippocampus-dependent memory tasks. (A) Object recognition task showing the percentage of exploration time directed toward the novel versus familiar object at 5 min, 30 min, and 24 hr after training. (B) Object location task assessing preference for the novel spatial location at 5 and 30 min after training. (C) Object-in-place task evaluating associative spatial recognition memory at 5 and 30 min after training. (D) Social recognition task measuring interaction time with familiar and novel social targets. (E) Object recency task assessing temporal order memory by comparing exploration of recent and old objects. (F) Contextual fear conditioning task showing freezing behavior during re-exposure to the conditioning chamber at 1 and 24 hr after training. (G) Y-maze spontaneous alternation task assessing spatial working memory; total arm entries and alternation performance are shown. Open circles indicate training sessions and closed circles indicate retrieval sessions (F, G). Sample sizes are shown below each bar. Data are presented as individual values with mean ± SEM. ^a^*P* < 0.05, ^aa^*P* < 0.01 *vs* familiar target or training; \**P* < 0.05, \*\**P* < 0.01 *vs* saline; ^#^*P* < 0.05, ^##^*P* < 0.01 *vs* Aβ.

In the object location task (**Figure 3B**), after a 5-min interval, the saline (*t* = 4.518, *P* = 0.0015), RLZ (*t* = 8.792, *P* < 0.0001), and Aβ +RLZ groups (*t* = 3.800, *P* = 0.0067) – but not the Aβ_1-42_ group (*t* = 0.409, *P* = 0.6936, paired t-test) – preferentially explored the novel location. RLZ injection improved Aβ-induced impairment in object location memory after 5 min. A similar pattern was observed 30 minutes later, with significantly higher exploration time in the novel location in the saline (*t* = 3.100, *P* = 0.0127), RLZ (*t* = 3.186, *P* = 0.0154), and Aβ+RLZ groups (*t* = 3.3958, *P* = 0.0055), but not in the Aβ_1-42_ group (*t* = 1.277, *P* = 0.2376, paired t-test). Aβ-induced impairment was also improved in the Aβ+RLZ group after 30 min. However, after 24 hours, no differences were detected among the groups.

In the object-in-place task (**Figure 3C**) the saline (*t* = 4.484, *P* = 0.0015), RLZ (*t* = 2.586, *P* = 0.0361), and Aβ +RLZ groups (*t* = 2.746, *P* = 0.0165) showed increased touch time of the exchanged objects after 5 min, while the Aβ_1-42_ group did not (*t* = 1.152, *P* = 0.2826, paired t-test). After 30 min, touch time of the exchanged objects increased in the saline (*t* = 3.160, *P* = 0.0116), RLZ (*t* =2.469, *P* = 0.0429), and Aβ+RLZ groups (*t* = 2.717, *P* = 0.0228), but not in the Aβ_1-42_ group (*t* = 1.862, *P* = 0.0996, paired t-test). Aβ-induced impairment in object association memory improved after 5- and 30-min intervals following RLZ injection. However, no significant differences were observed after 24 hr.

In the social recognition task (**Figure 3D**) the saline (*t* = 3.550, *P* = 0.0062), RLZ (*t* = 2.932, *P* = 0.0220), and Aβ+RLZ groups (*t* = 2.506, *P* = 0.0406) interacted more with the novel social target after a 5-min test interval, though the Aβ_1-42_ group did not (*t* = 0.095, *P* = 0.9263, paired t-test). In the object recency task (**Figure 3E**), the time spent touching the old object was significantly longer in the saline (*t* = 4.649, *P* = 0.0012), RLZ (*t* = 2.524, *P* = 0.0396), and Aβ+RLZ groups (*t* = 2.451, *P* = 0.0440), after the second sampling phase, but not in the Aβ_1-42_ group (*t* = 0.312, *P* = 0.7632, paired t-test).

In the fear conditioning test (**Figure 3F**), after 1 hr post-training, a three-way ANOVA revealed main effects of training (*F*_1,62_ = 155.970, *P* < 0.0001), Aβ (*F*_1,62_ = 12.515, *P* = 0.0008) and riluzole (*F*_1,62_ = 7.882, *P* = 0.0067). Freezing time increased in the saline, RLZ, and Aβ+RLZ groups (all; *p* < 0.0001 *vs* training) but not in the Aβ group (*p* = 0.1320 *vs* training). Aβ impaired contextual memory (Aβ *vs* saline; *p* < 0.0001), which improved after riluzole injection (Aβ *vs* Aβ+RLZ; *p* = 0.0016). After 24 hr, a three-way ANOVA revealed main effects of training (*F*_1,62_ = 146.670, *P* < 0.0001) and riluzole (*F*_1,62_ = 7.104, *P* = 0.0098) but no significant Aβ effect (*F*_1,62_ = 3.491, *P* = 0.0664). Freezing time increased in all groups (all; *P* < 0.0001 *vs* training) including the Aβ group (*P* = 0.0111 *vs* training), and the Aβ group showed partial impairment alleviated by RLZ injection (Aβ *vs* Aβ+RLZ; *P* = 0.0068).

In the Y-maze task (**Figure 3G**), arm entries tended to decrease after 24 hr, though this change was not statistically significant (saline: *t* = 1.397, *P* = 0.1959; RLZ: *t* = 1.795, *P* = 0.1158; Aβ: *t* = 2.121, *p* = 0.0667; Aβ+RLZ: *t* = 1.140, *P* = 0.2920, paired t-test). Training increased the alternation ratio in the saline (*t* = 3.052, *P* = 0.0138) and RLZ groups (*t* = 2.829, *P* = 0.0254) but not in the Aβ (*t* = 2.289, *P* = 0.0514) and Aβ +RLZ groups after 24 hr (*t* = 1.811, *P* = 0.1131, paired t-test). Two-way ANOVA revealed main effects of Aβ (*F*_1,31_ = 7.985, *P* = 0.0082) and riluzole (*F*_1,31_ = 8.969, *P* = 0.0054), as well as a significant interaction (*F*_1,31_ = 7.670, *P* = 0.0094). Aβ-induced impairment in spatial working memory (Aβ *vs* saline; *P* = 0.0015) improved in the Aβ+RLZ group (Aβ *vs* Aβ+RLZ; *P* = 0.0020). Together, these results demonstrate that riluzole broadly rescues Aβ_1-42_ oligomer–induced impairments across multiple hippocampus-dependent memory domains, without affecting exploratory activity or motivation.

### The effect of riluzole on Aβ_1-42_ oligomer deposition and clearance by astrocytes and microglia

First, we examined the effects of riluzole on the deposition of Aβ_1-42_ oligomers in neurons and their clearance by astrocytes and microglia. Aβ/NeuN^+^ cells increased markedly in the Aβ group and decreased in the Aβ+RLZ group (**Figure 4A**, Aβ: *F_1,44_* = 1236.140, *P* < 0.0001; riluzole: *F_1,44_* = 225.432, *P* < 0.0001; interaction: *F_1,44_* = 215.664, *P* < 0.0001). Next, the number of Aβ/GFAP^+^ cells were increased in the Aβ group and increased further in the Aβ+RLZ group (**Figure 4B**, Aβ: *F_1,54_* = 356.810, *P* < 0.0001; riluzole: *F_1,54_* =30.293, *P* < 0.0001; interaction: *F_1,54_* =30.293, *P* < 0.0001). Super-resolution microscopy of z-stack (4 µm) images further confirmed internalized Aβ in astrocytes (**Figure S1)**. Similarly, the number of Aβ/Iba1^+^ cells were increased in the Aβ group and was further elevated in the Aβ+RLZ group (**Figure 4C**, Aβ: *F_1,38_* = 279.172, *P* < 0.0001; riluzole: *F_1,38_* = 76.638, *P* < 0.0001; interaction: *F_1,38_* = 76.638, *P* < 0.0001). Therefore, riluzole reduced the number of Aβ-positive neurons while enhancing Aβ binding by astrocytes and microglia.

**Figure 4.**
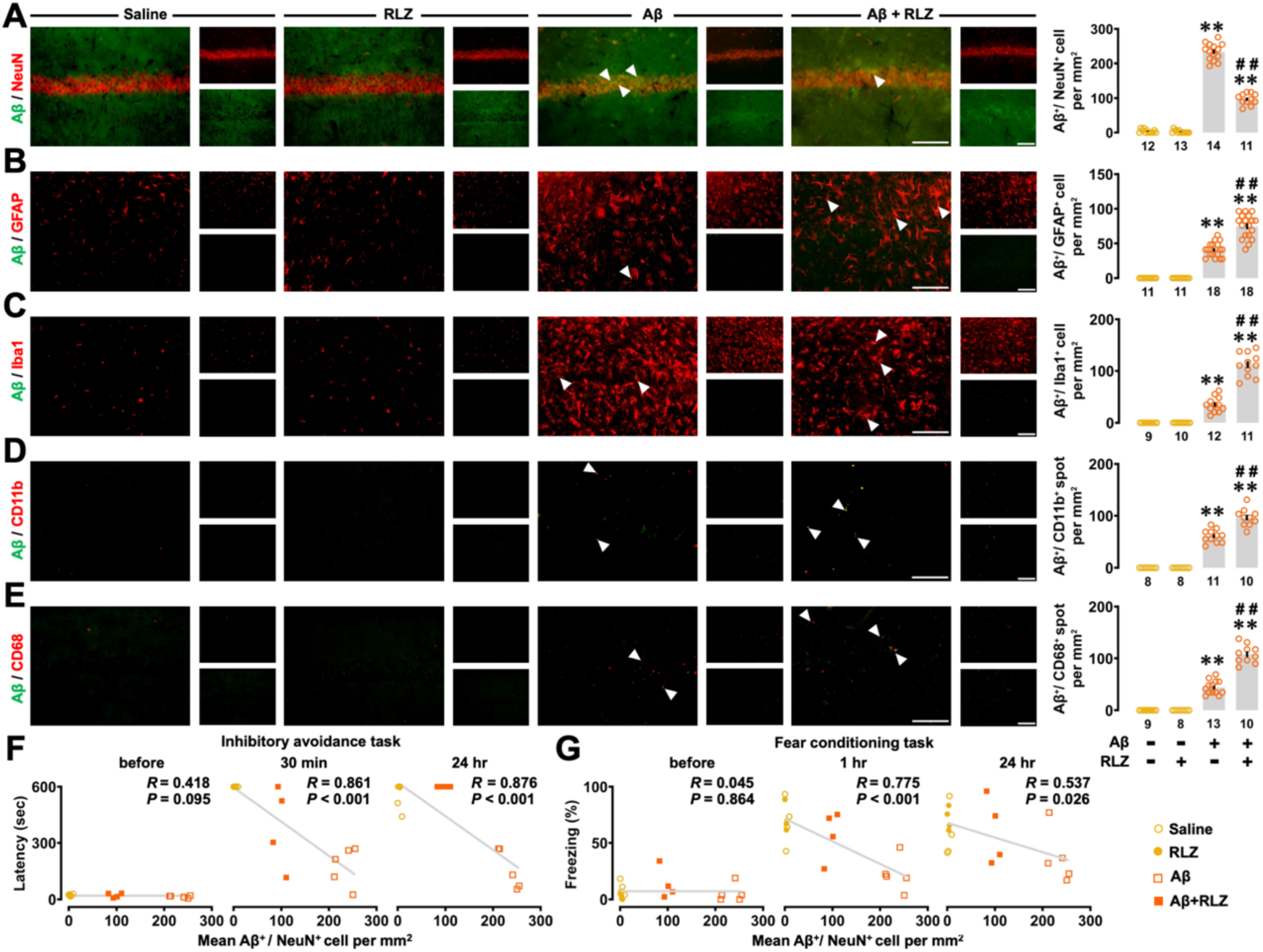
Riluzole alters the cellular distribution of Aβ_1-42_ oligomers in the dorsal CA1 region. (A) Representative fluorescence images and quantification of lecanemab-positive Aβ within NeuN⁺ neurons. The number of Aβ⁺ neurons was reduced in the Aβ+RLZ group. (B, C) Representative images and quantification of lecanemab-positive Aβ associated with GFAP⁺ astrocytes (B) and Iba1⁺ microglia (C). The number of Aβ-associated astrocytes and microglia were increased in the Aβ+RLZ group. (D, E) Representative images and quantification of colocalized Aβ/CD11b (D) and Aβ/CD68 (E) puncta per mm². (F, G) Correlation analyses between learning performance and neuronal Aβ burden. Learning performance after training, but not before, was negatively correlated with mean number of Aβ/NeuN^+^ neurons in the IA task (F) and contextual fear conditioning (G). Each dot represents an individual animal. Arrowheads indicate double-positive cells or colocalized puncta. Scale bars: 100 μm. Sample sizes are shown below each bar. Data are presented as individual values with mean ± SEM. \*\**P* < 0.01 *vs* saline; ^##^*P* < 0.01 *vs* Aβ.

Next, we examined the effect of riluzole on Aβ recognition by CD11b and degradation by CD68. Colocalized Aβ/CD11b spots increased in the Aβ group and increased further in the Aβ+RLZ group (**Figure 4D**, Aβ: *F_1,33_* = 434.753, *P* < 0.0001; riluzole: *F_1,33_* = 22.421, *P* < 0.0001; interaction: *F_1,33_* = 22.421, *P* < 0.0001). Similarly, colocalized Aβ/CD68 spots increased in the Aβ group, and this increase was further elevated in the Aβ+RLZ group (**Figure 4E**, Aβ: *F_1,36_* =404.181, *P* < 0.0001; riluzole: *F_1,36_* = 71.911, *p* < 0.0001; interaction: *F_1,36_* = 71.911, *P* < 0.0001). These data highlight that riluzole enhances CD11b-mediated Aβ recognition and lysosome-mediated Aβ degradation by microglia.

To further establish a brain–behavior link, we examined correlations between contextual learning performance and the neuronal Aβ burden, quantified as the mean number of Aβ/NeuN^+^ cells in individual animals. No significant correlation was detected prior to the IA task (*R* = −0.418, *P* = 0.0951). In contrast, a strong negative correlation emerged between the number of Aβ/NeuN^+^ cells and IA performance measured at both 30 min (*R* = −0.861, *P* < 0.0001) and 24 hr (*R* = −0.876, *P* < 0.0001) after training (**Figure 4F**).

A similar relationship was observed in the contextual fear conditioning paradigm (**Figure 4G**). Before shock exposure, freezing behavior did not correlate with neuronal Aβ burden (*R* = −0.045, *P* = 0.8639). However, significant negative correlations were detected between the number of Aβ/NeuN^+^ cells and freezing behavior at 1 h (*R* = −0.775, *P* = 0.0003) and 24 h (*R* = −0.537, *P* = 0.0262) after conditioning. Together, these results indicate that reductions in neuronal Aβ burden—consistent with enhanced glial-mediated Aβ clearance—are tightly associated with improved hippocampus-dependent learning and memory performance.

### Effect of riluzole on Aβ_1-42_ oligomer-induced neuronal apoptosis, spine loss and alterations in spine morphology

We examined the effects of riluzole on Aβ_1-42_ oligomer-induced neuronal apoptosis and dendritic degeneration. First, Aβ_1-42_ oligomers reduced the number of NeuN^+^ cells (Aβ *vs* saline, *P* < 0.0001); but this effect was reversed in the Aβ+RLZ group (**Figure 5A**, Aβ: *F_1,49_* =31.761, *P* < 0.0001; riluzole: *F_1,49_* =38.354, *P* < 0.0001; interaction: *F_1,49_* = 27.513, *P* < 0.0001). Similarly, the number of cleaved caspase-3/NeuN^+^ cells was higher in the Aβ group and lower in the Aβ+RLZ group (**Figure 5B**, Aβ: *F_1,39_* = 169.243, *P* < 0.0001; riluzole: *F_1,39_* = 95.698, *P* < 0.0001; interaction: *F_1,39_* = 103.523, *P* < 0.0001). Conversely, the Aβ-induced reduction in MAP2 intensity was reversed in the Aβ + RLZ group (**Figure 5C**, Aβ: *F_1,60_* = 212.960, *P* < 0.0001; riluzole: *F_1,60_* = 122.056, *P* < 0.0001; interaction: *F_1,60_* = 116.174, *P* < 0.0001). These data suggest that riluzole mitigates Aβ_1-42_ oligomer-induced neuronal apoptosis and dendritic degeneration.

**Figure 5.**
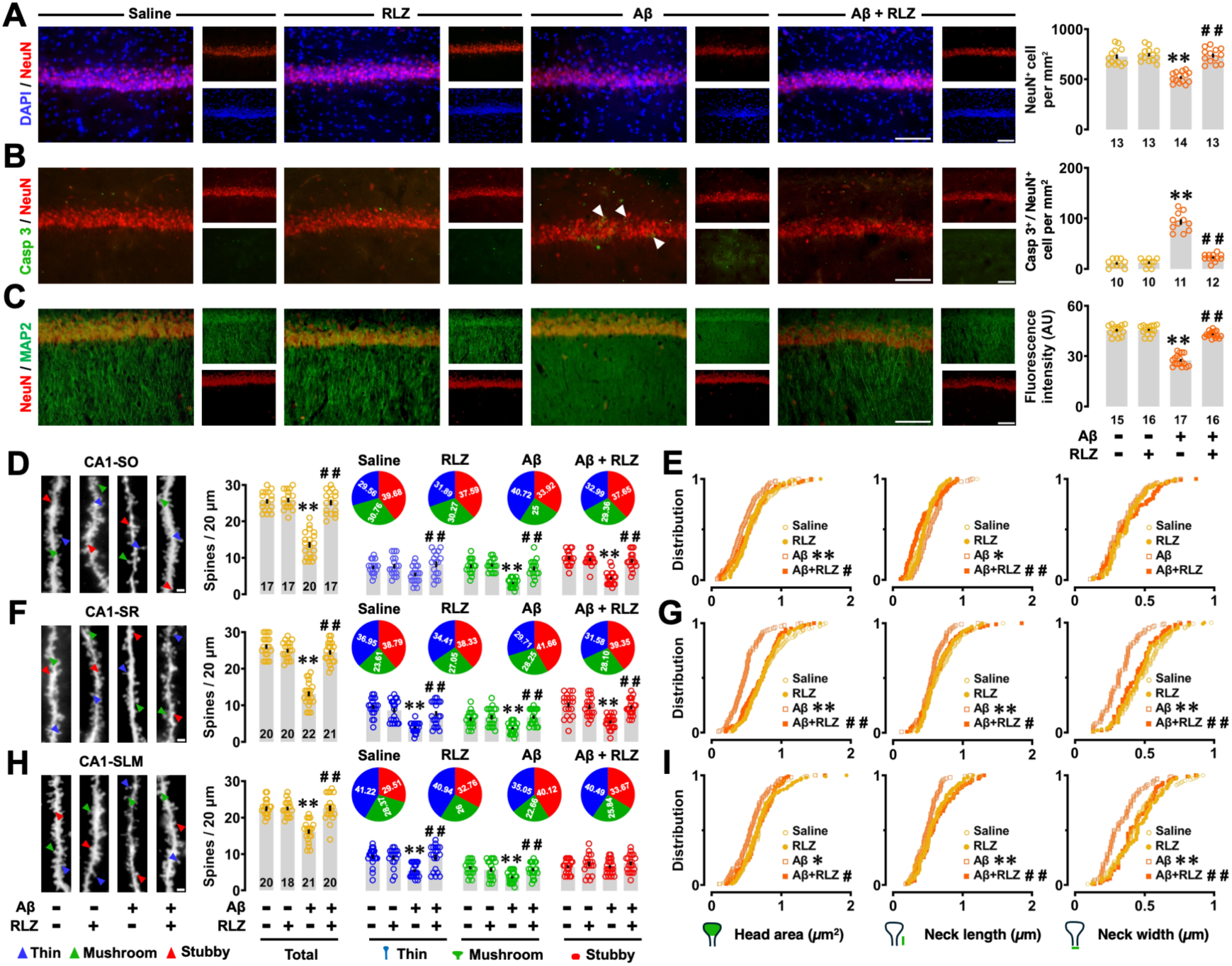
Effects of riluzole on neuronal survival, dendritic integrity, and spine morphology in the dorsal CA1 region. (A, B) Representative fluorescence images and quantification of NeuN⁺ cells (A) and cleaved caspase-3/NeuN⁺ cells (B) in the dorsal CA1. (C) Representative images and quantification of MAP2 fluorescence intensity in the stratum radiatum of the dorsal CA1. (D, F, H) Representative Golgi-stained images and quantification of dendritic spine density (spines per 20 μm) in the stratum oriens (D), stratum radiatum (F), and stratum lacunosum-moleculare (H). Pie charts indicate the proportion of thin, mushroom, and stubby spines in each group. (E, G, I) Cumulative distributions of spine head area, neck length, and neck width for individual spines in the stratum oriens (E), stratum radiatum (G), and stratum lacunosum-moleculare (I). Arrowheads indicate double-positive cells. Images in (D, F, H) show Golgi-stained dendrites pseudocolored in gray. Colored arrowheads indicate spine types. Scale bars: 100 μm (A–C), 2 μm (D, F, H). Sample sizes (number of sections, regions of interest, or dendrites) are shown below each bar. Data are presented as individual values with mean ± SEM. \**P* < 0.05, \*\**P* < 0.01 *vs* saline; ^##^*P* < 0.01 *vs* Aβ.

Next, in the stratum oriens (**Figure 5D**), the loss of total spines, as well as mushroom and stubby spines, induced by Aβ was reversed in the Aβ + RLZ group. This was accompanied by the reversal of smaller Aβ-induced spine heads and shorter necks of individual spines in the Aβ+RLZ group (**Figure 5E**). In the stratum radiatum (**Figure 5F**), Aβ reduced the total spine density and each spine type. These effects were reversed in the Aβ + RLZ group. Furthermore, riluzole restored the smaller heads, shorter necks and narrower necks induced by Aβ (**Figure 5G**). In the stratum lacunosum-moleculare (**Figure 5H**), Aβ induced a loss of total spines, as well as thin and mushroom spines. These were recovered in the Aβ + RLZ group. In the Aβ+RLZ group, riluzole reversed the smaller heads, shorter necks, and narrower necks induced by Aβ (**Figure 5I**). Overall, our findings demonstrate that riluzole reverses the effects of Aβ_1-42_ oligomers on spine loss and alterations in spine morphology (see **Table S2** for statistical results).

### Effect of riluzole on Aβ_1-42_ oligomer-induced complement-mediated synaptic elimination by microglia

To determine whether riluzole modulates synaptic integrity under Aβ oligomer exposure, we first examined complement-associated synaptic alterations in the dorsal CA1 region. To investigate how riluzole rescued Aβ-induced synaptic loss, we quantified the number of colocalized vGlut1/PSD95 puncta. In the stratum radiatum, the number of synaptic puncta decreased in the Aβ group, but remained stable in the Aβ+RLZ group (**Figure 6A**, Aβ: *F_1,35_* = 27.207, *P* < 0.0001; riluzole: *F_1,35_* = 7.560, *P* = 0.0094; interaction: *F_1,35_* = 17.638, *P* = 0.0002). Similar changes were also observed in the stratum lacunosum-moleculare (**Figure S2A**). However, the reduction in synaptic puncta was attributable to the loss of the postsynaptic protein PSD95 rather than the presynaptic marker vGlut1 (**Figure S2B**; **Table S3** for statistical results).

**Figure 6.**
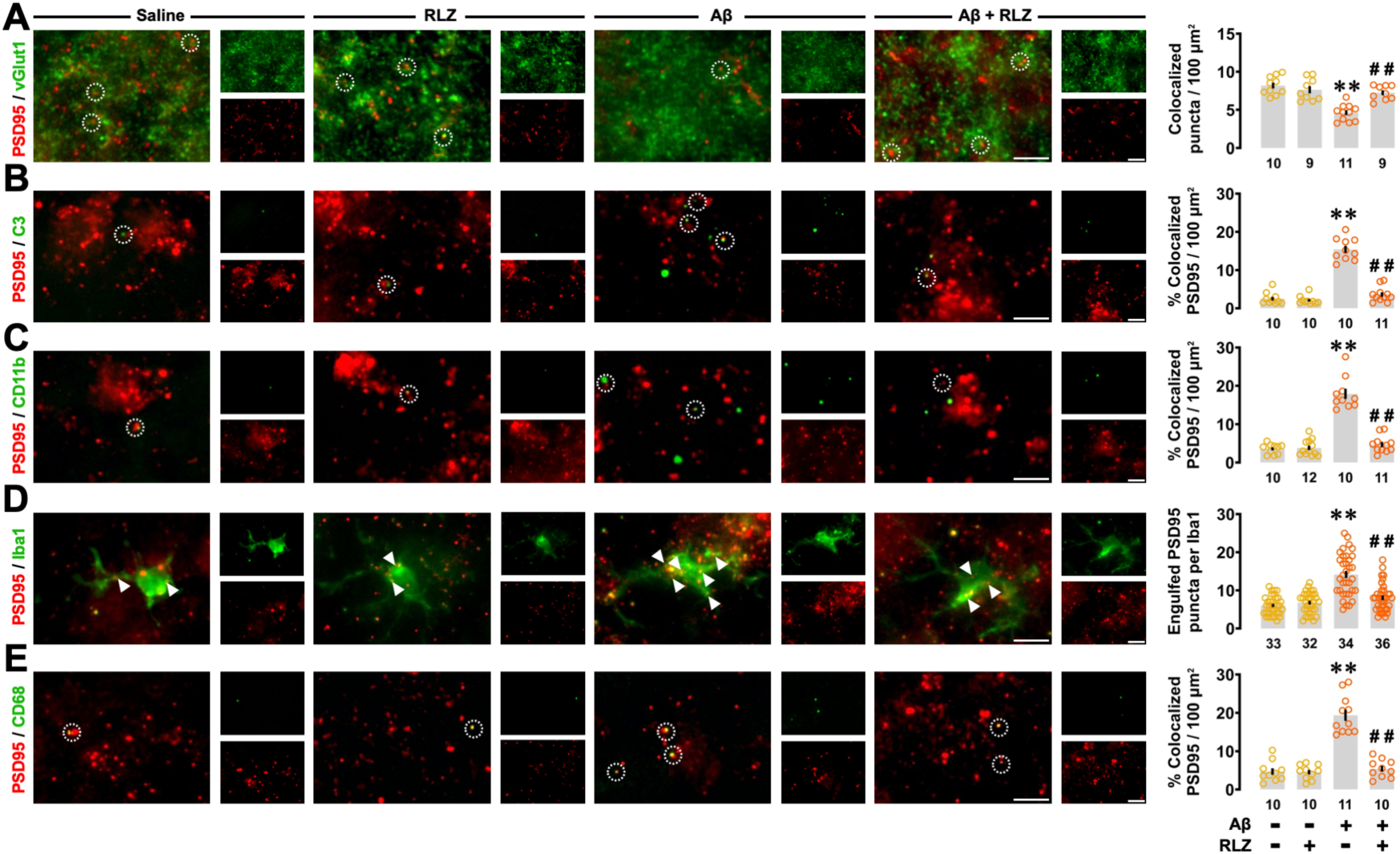
Effects of riluzole on synaptic markers and microglial-associated synaptic puncta in the dorsal CA1 region. (A) Representative fluorescence images and quantification of colocalized vGlut1/PSD95 puncta per 100 μm² in the stratum radiatum of the dorsal CA1. (B, C) Representative images and quantification of colocalized C3/PSD95 (B) and CD11b/PSD95 (C) puncta per 100 μm². (D) Representative images and quantification of PSD95 puncta engulfed by Iba1⁺ microglia. (E) Representative images and quantification of colocalized CD68/PSD95 puncta per 100 μm². Dotted circles indicate colocalized puncta and arrowheads indicate engulfed PSD95 puncta. Scale bars: 5 μm. Sample sizes (number of regions of interest or cells) are shown below each bar. Data are presented as individual values with mean ± SEM. \*\**P* < 0.01 *vs* saline; ^#^*P* < 0.05, ^##^*P* < 0.01 *vs* Aβ.

Next, we examined complement-mediated synaptic engulfment by microglia. C3/PSD95 colocalization increased in the Aβ group but remained at control levels in the Aβ+RLZ group (**Figure 6B**, Aβ: *F_1,37_* = 130.757, *P* < 0.0001; riluzole: *F_1,37_* = 94.518, *P* < 0.0001; interaction: *F_1,37_* = 82.087, *P* < 0.0001). CD11b/PSD95 colocalization was also increased in the Aβ group and reduced in the Aβ+RLZ group (**Figure 6C**, Aβ: *F_1,39_* = 88.627, *P* < 0.0001; riluzole: *F_1,39_* = 63.999, *P* < 0.0001; interaction: *F_1,39_* = 69.316, *p* < 0.0001). Consistently, the number of PSD95 puncta within microglia was elevated in the Aβ group and restored in the Aβ+RLZ group (**Figure 6D**, Aβ: *F_1,31_* = 51.139, *P* < 0.0001; riluzole: *F_1,31_* = 16.538, *P* < 0.0001; interaction: *F_1,31_* = 27.148, *P* < 0.0001). CD68/PSD95 colocalization showed a similar pattern (**Figure 6E**, Aβ: *F_1,37_* = 57.974, *P* < 0.0001; riluzole: *F_1,37_* = 47.782, *P* < 0.0001; interaction: *F_1,37_* = 45.639, *P* < 0.0001). These results indicate that riluzole attenuates Aβ_1-42_ oligomer–induced synaptic loss in parallel with reduced complement-associated microglial synaptic elimination.

### Effect of riluzole on Aβ_1-42_ oligomer–induced astrocyte recruitment, morphology, and marker expression

Given the close involvement of astrocytes in complement signalling and synaptic regulation, we next assessed astrocytic responses in the same hippocampal region. Two-way ANOVA revealed that Aβ_1-42_ oligomers triggered robust recruitment of GFAP⁺ astrocytes, which was modestly reduced but persisted in the Aβ+RLZ group (**Figure 7A**, Aβ: *F_1,66_* = 248.292, *P* < 0.0001; riluzole: *F_1,66_* = 14.009, *P* = 0.0004; interaction: *F_1,66_* = 19.323, *p* < 0.0001).

**Figure 7.**
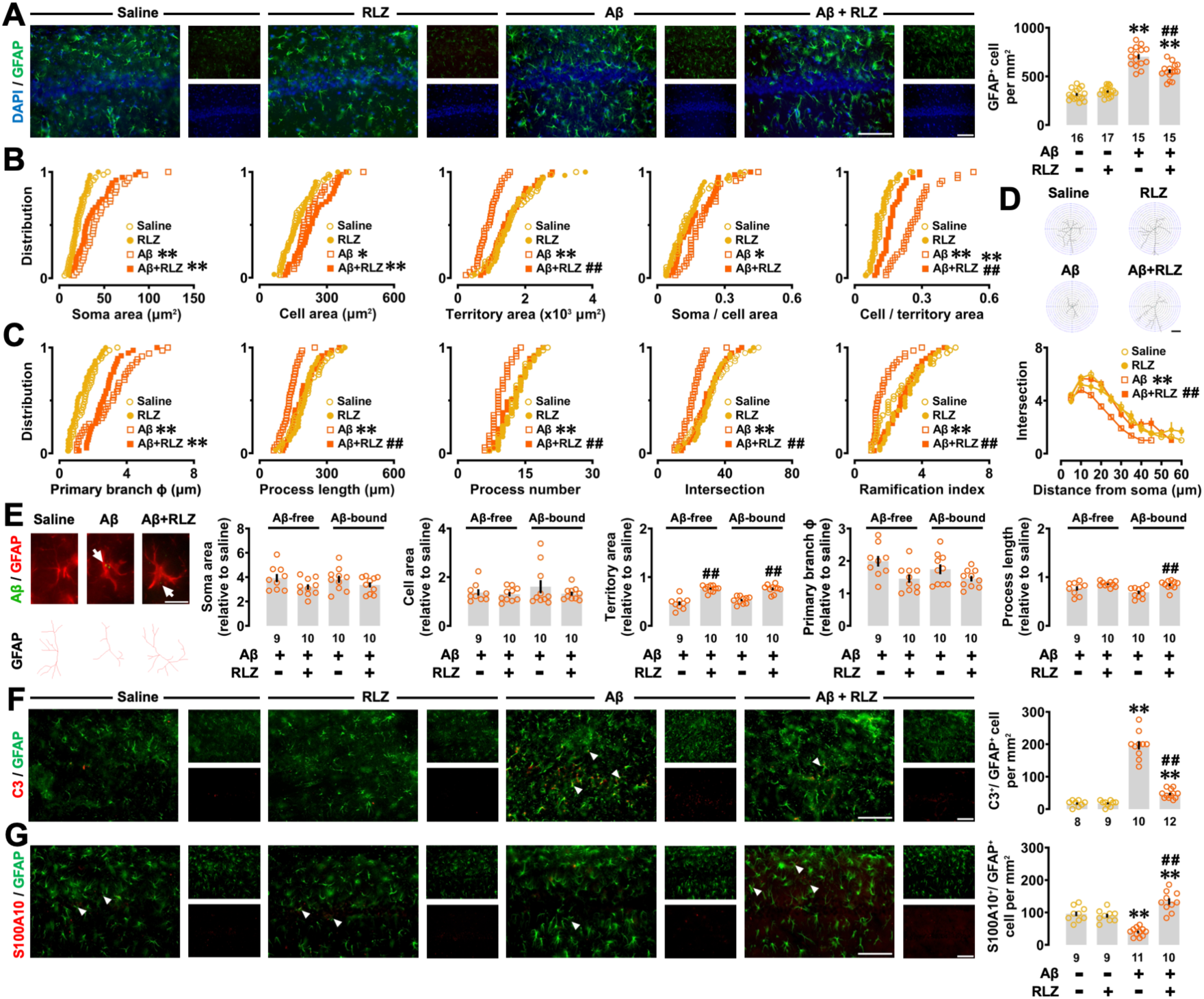
Effects of riluzole on astrocyte number, morphology, and polarization in the dorsal CA1 region. (A) Representative fluorescence images and quantification of GFAP⁺ cells per mm² in the dorsal CA1 region. (B) Quantification of size- and shape-related astrocytic parameters, including soma area, cell area, territory area, and their ratios. (C) Quantification of branching-related parameters, including primary branch thickness, process length, process number, intersections, and ramification index. (D) Representative reconstructed astrocyte skeletons (top) and Sholl analysis (bottom). Intersections were quantified at 5-µm intervals from the soma. (E) Representative fluorescence images of lecanemab-positive astrocytes (top) and corresponding reconstructed skeletons (bottom), with morphological parameters analyzed separately for Aβ-free and Aβ-bound astrocytes. (F, G) Representative fluorescence images and quantification of C3/GFAP⁺ (F) and S100A10/GFAP⁺ (G) cells per mm². Arrowheads indicate double-positive cells and arrows indicate Aβ₁₋₄₂ oligomers. Scale bars: 100 μm (A, F, G) and 20 μm (D, E). Sample sizes (number of slices or cells) are shown below each bar. Data are presented as individual values with mean ± SEM. \**P* < 0.05, \*\**P* < 0.01 *vs* saline; ^##^*P* < 0.01 *vs* Aβ.

Regarding morphology, soma and cell area increased in both the Aβ and Aβ+RLZ groups (**Figure 7B**). Territory area decreased in the Aβ group but increased in the Aβ+RLZ group, while changes in soma-to-cell and cell-to-territory area ratios were partially normalized. Analysis of branching complexity showed increased primary branch thickness in both groups (**Figure 7C**). In contrast, Aβ-induced reductions in total process length, process number, intersections, and ramification index were reversed in the Aβ+RLZ group. Sholl analysis confirmed recovery of branching complexity (**Figure 7D**).

Morphological comparison between Aβ-free and Aβ-bound astrocytes revealed no differences in soma area or cell area; however, territory area (Aβ oligomers: *F_1,35_* = 1.081, *P* = 0.3056; riluzole: *F_1,35_* = 85.201, *P* < 0.0001; interaction: *F_1,35_* = 1.474, *P* = 0.2328) and process length (Aβ oligomers: *F_1,35_* = 2.075, *P* = 0.1587; riluzole: *F_1,35_* = 14.589, *P* = 0.0005; interaction: *F_1,35_* = 0.948, *P* = 0.3370) were increased in the Aβ+RLZ group regardless of Aβ binding (**Figure 7E**). Finally, C3⁺/GFAP⁺ astrocytes were increased in the Aβ group and significantly reduced in the Aβ+RLZ group, whereas S100A10⁺/GFAP⁺ astrocytes showed the opposite pattern (**Figure 7F**, Aβ: *F_1,35_* = 230.772, *P* < 0.0001; riluzole: *F_1,35_* = 107.198, *P* < 0.0001; interaction: *F_1,35_* = 107.895, *P* < 0.0001. **Figure 7G**, Aβ: *F_1,35_* = 0.648, *P* = 0.4264; riluzole: *F_1,35_* = 37.216, *P* < 0.0001; interaction: *F_1,35_* = 48.352, *P* < 0.0001).

Together, these findings show that riluzole preserves astrocyte recruitment while normalizing astrocytic morphology and marker expression under Aβ conditions.

### Effect of riluzole on Aβ_1-42_ oligomer–induced microglial recruitment, morphology, and functional marker expression

We then examined whether microglial structural and molecular responses were similarly modulated by riluzole. Aβ_1-42_ oligomers significantly increased Iba1⁺ microglial density, which was modestly reduced but persisted in the Aβ+RLZ group (**Figure 8A**, Aβ: *F_1,62_* = 208.608, *P* < 0.0001; riluzole: *F_1,62_* = 6.611, *P* = 0.0125; interaction: *F_1,62_* = 6.100, *P* = 0.0163).

**Figure 8.**
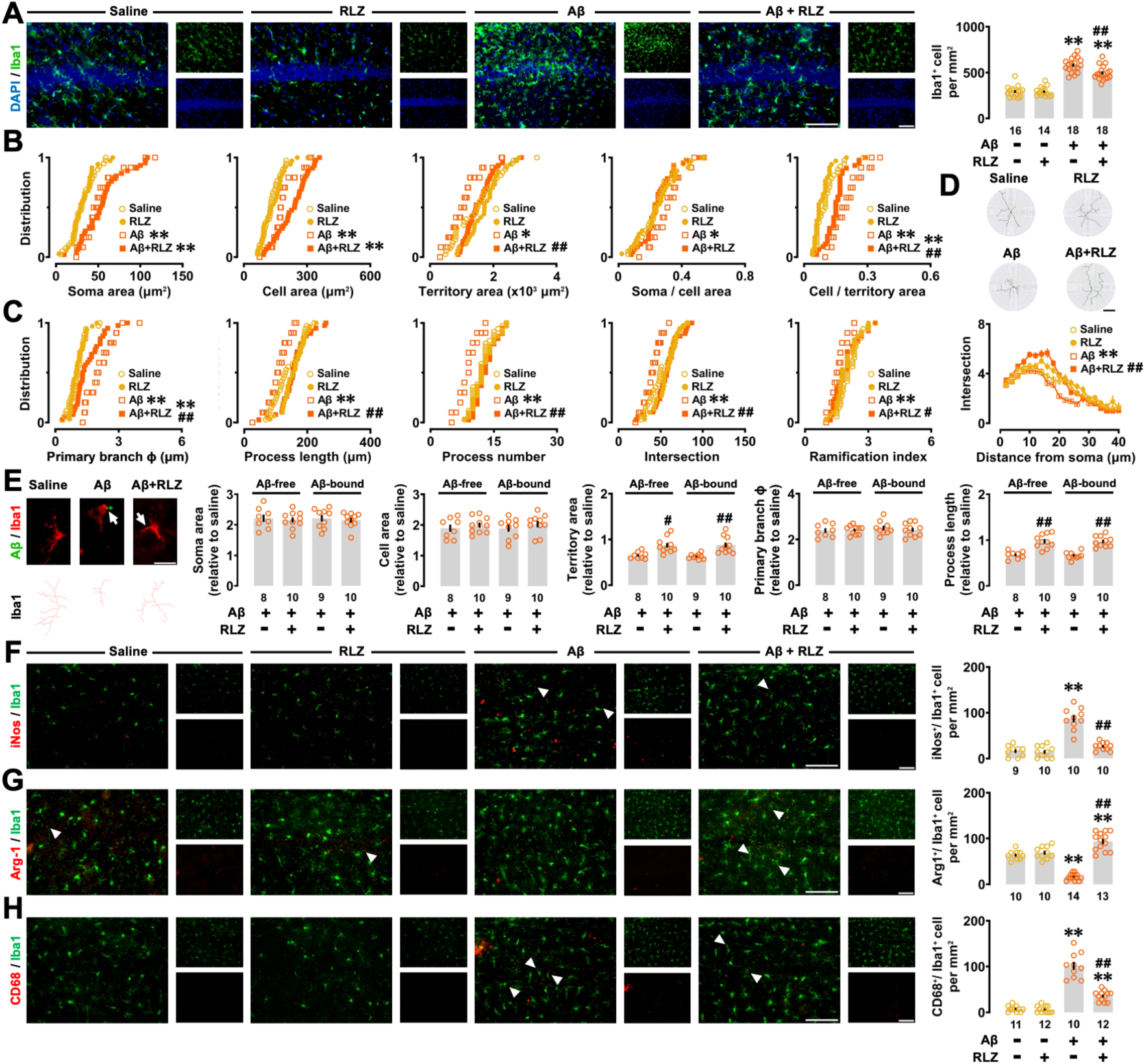
Effects of riluzole on microglial number, morphology, and marker expression in the dorsal CA1 region. (A) Representative fluorescence images and quantification of Iba1⁺ cells per mm² in the dorsal CA1 region. (B) Quantification of size- and shape-related microglial parameters, including soma area, cell area, territory area, and their ratios. (C) Quantification of branching-related parameters, including primary branch thickness, process length, process number, intersections, and ramification index. (D) Representative reconstructed microglial skeletons (top) and Sholl analysis (bottom). Intersections were quantified at 2-µm intervals from the soma. (E) Representative fluorescence images of lecanemab-positive microglia (top) and corresponding reconstructed skeletons (bottom), with morphological parameters analyzed separately for Aβ-free and Aβ-bound microglia. (F, G, H) Representative fluorescence images and quantification of iNOS/Iba1⁺ (F), Arg1/Iba1⁺ (G), and CD68/Iba1⁺ (H) cells per mm². Arrowheads indicate double-positive cells and arrows indicate Aβ_1-42_ oligomers. Scale bars: 100 μm (A, F–H) and 20 μm (D, E). Sample sizes (number of slices or cells) are shown below each bar. Data are presented as individual values with mean ± SEM. \**P* < 0.05, \*\**P* < 0.01 *vs* saline; ^#^*P* < 0.05, ^##^*P* < 0.01 *vs* Aβ.

Morphological analysis revealed increased soma and cell area in both groups (**Figure 8B**). Aβ-induced reductions in territory size and increases in cell-to-territory area ratio were partially normalized in the Aβ+RLZ group. Branching analysis showed that reductions in process length, process number, intersections, and ramification index in the Aβ group were reversed by riluzole (**Figure 8C, D**).

Analysis of Aβ-free and Aβ-bound microglia showed increased territory area (Aβ oligomers: *F_1,33_* = 0.040, *P* = 0.8432; riluzole: *F_1,33_* = 26.458, *P* < 0.0001; interaction: *F_1,33_* = 0.082, *P* = 0.7760) and process length (Aβ oligomers: *F_1,33_* = 0.058, *P* = 0.8104; riluzole: *F_1,33_* = 64.549, *P* < 0.0001; interaction: *F_1,33_* = 0.026, *P* = 0.8720) in the Aβ+RLZ group independent of Aβ binding (**Figure 8E**). Consistent with these structural changes, riluzole reduced the proportion of iNOS⁺/Iba1⁺ microglia while increasing Arg1⁺/Iba1⁺ cells and decreasing CD68⁺/Iba1⁺ microglia (**Figure 8F**, Aβ: *F_1,35_* = 68.148, *P* < 0.0001; riluzole: *F_1,35_* = 38.833, *P* < 0.0001; interaction: *F_1,35_* = 34.924, *P* < 0.0001. **Figure 8G**, Aβ: *F_1,43_* = 8.117, *P* = 0.0067; riluzole: *F_1,43_* = 106.717, *P* < 0.0001; interaction: *F_1,43_* = 80.118, *P* < 0.0001. **Figure 8H**, Aβ: *F_1,41_* = 187.003, *P* < 0.0001; riluzole: *F_1,41_* = 55.255, *P* < 0.0001; interaction: *F_1,41_* = 51.355, *P* < 0.0001). Additionally, **Figure S3** shows the percentage of different morphological types of microglia in each group (see **Table S4** for statistical results). These data indicate that riluzole modulates microglial morphology and functional marker expression in parallel with reduced synaptic engulfment.

## Discussion

### Riluzole improves Aβ_1-42_ oligomer–induced memory impairment by enhancing glial-mediated Aβ handling

In the present study, we demonstrate that riluzole ameliorates Aβ_1-42_ oligomer–induced cognitive impairment and hippocampal pathology by enhancing glial-mediated handling of Aβ. Across individual animals, hippocampus-dependent learning performance was strongly and negatively correlated with neuronal Aβ burden, providing a direct behavioral link between cellular Aβ distribution and memory outcome. Importantly, riluzole did not suppress glial recruitment; instead, it reshaped astrocytic and microglial responses toward functional states associated with efficient Aβ uptake, degradation, and synaptic preservation.

Soluble Aβ oligomers disrupt glutamatergic signaling (Tu et al., 2014) and induce neuronal hyperexcitability (Dickerson et al., 2005; Zott et al., 2019), ultimately leading to cognitive impairment (Lei et al., 2016; Yang et al., 2021). Aβ-induced neurotoxicity is tightly coupled to glutamatergic dysfunction (Li et al., 2011), which in turn promotes amyloid precursor protein processing and further Aβ release (Bordji et al., 2010), creating a self-amplifying pathological loop. Interventions that stabilize glutamatergic signaling may therefore indirectly facilitate Aβ clearance. In this context, the cognitive benefits of riluzole observed here are most parsimoniously explained by enhanced removal and redistribution of Aβ rather than by reduced amyloid production.

Consistent with our findings, riluzole improves cognitive performance and reduces amyloid burden in both transgenic(Hunsberger et al., 2015; Okamoto et al., 2018; Hascup et al., 2021) and Aβ injection–based models (Yang et al., 2021) of AD. Because riluzole does not alter amyloidogenic APP (amyloid precursor protein) processing (Okamoto et al., 2018), these effects have been attributed to enhanced Aβ clearance mechanisms. Our data extend these observations by demonstrating that riluzole shifts Aβ from neurons toward glial compartments in vivo and that this redistribution is closely associated with improved learning and memory at the individual-animal level.

Both astrocytes and microglia contribute to Aβ handling, although their relative roles remain debated. While microglial uptake of Aβ often exceeds that of astrocytes (Serrano-Pozo et al., 2013), coordinated astrocyte–microglia interactions promote more efficient Aβ clearance than either cell type acting alone (Rostami et al., 2021). Our findings support this cooperative model, indicating that riluzole enhances complementary glial functions rather than selectively activating a single cell population.

### Riluzole preserves synapses and neuronal structure by limiting excitotoxicity and complement-mediated pruning

In parallel with enhanced glial Aβ handling, riluzole attenuated Aβ oligomer–induced neuronal apoptosis, dendritic degeneration, and dendritic spine loss across the dorsal CA1 sublayers. Aβ-induced excitotoxicity drives excessive Ca²⁺ influx and mitochondrial dysfunction, culminating in neuronal death (Calvo-Rodriguez et al., 2020). Riluzole limits this process by reducing presynaptic glutamate release (Hascup et al., 2021) and enhancing glutamate uptake (Fumagalli et al., 2008), thereby mitigating excitotoxic stress. Consistent with prior reports across multiple disease models (Wu et al., 2013; Kyllo et al., 2024; Shen et al., 2025), our findings indicate that riluzole effectively protects neurons from Aβ-induced apoptotic degeneration.

Dendritic degeneration and spine loss are strongly associated with cognitive dysfunction in AD (Knobloch & Mansuy, 2008). Because dendritic architecture determines neuronal integrative properties, dendritic simplification contributes to network hyperexcitability (Šišková et al., 2014) and impaired information processing (Koh et al., 2010). The preservation of dendritic structure observed following riluzole treatment therefore provides a structural substrate for the recovery of hippocampus-dependent memory.

Aβ oligomers induce robust dendritic spine loss (Lacor et al., 2004; Wei et al., 2010), often accompanied by immature or aberrant spine morphologies (Lacor et al., 2007; Patnaik et al., 2020). In contrast, riluzole preserved spine density and structural integrity. These effects are consistent with previous observations that riluzole enhances synaptic organization (Pereira et al., 2017) and rescues Aβ-induced deficits in synaptic plasticity (Yang et al., 2021). Together, these findings support a model in which riluzole stabilizes synaptic structure and function under conditions of Aβ challenge.

Beyond direct excitotoxic mechanisms in AD, synapse loss is increasingly attributed to aberrant complement-mediated pruning (Tu et al., 2014). Soluble Aβ oligomers activate the classical complement cascade, leading to C3-dependent microglial engulfment of synapses, particularly postsynaptic elements (Almeida et al., 2005; Hong et al., 2016). This process closely coincides with dendritic spine loss and correlates strongly with cognitive decline.

Importantly, complement signaling also contributes to Aβ clearance, indicating that pathological synaptic pruning and beneficial amyloid removal are mediated by overlapping but dissociable pathways (Lv et al., 2024). In the present study, riluzole reduced C3- and CR3-associated synaptic engulfment and limited CD68-positive synapse internalization without suppressing microglial recruitment. These findings suggest that riluzole constrains aberrant complement-dependent synaptic elimination while preserving microglial phagocytic engagement.

Thus, riluzole appears to uncouple complement-mediated Aβ clearance from pathological synapse loss, preserving synaptic integrity and hippocampal circuit function while allowing effective amyloid handling.

### Coordinated astrocyte–microglia remodeling supports amyloid clearance and circuit resilience

A key finding of this study is that riluzole preserved astrocytic and microglial recruitment in response to Aβ oligomers while promoting functional and morphological states associated with neuroprotection. This contrasts with models in which glial activation is broadly suppressed and underscores the importance of glial functional state rather than glial presence per se.

Astrocyte accumulation around amyloid pathology is increasingly recognized as a protective response. Genetic disruption of astrocytic structural proteins exacerbates amyloid deposition and reduces astrocyte branching and plaque coverage (Kraft et al., 2013), highlighting the importance of astrocyte recruitment and physical engagement with amyloid pathology. In our study, riluzole sustained astrocyte recruitment while reducing features associated with neurotoxic reactivity.

Reactive A1 astrocytes, characterized by elevated C3 expression, lose key homeostatic functions, including aquaporin-4 polarization required for glymphatic Aβ clearance (Liddelow et al., 2017; Feng et al., 2023), and can impair microglial Aβ phagocytosis (Lian et al., 2016). In contrast, neuroprotective astrocyte states secrete trophic factors and anti-inflammatory mediators that support neuronal survival and promote amyloid clearance (Mizuta et al., 2001; El-Khatib et al., 2024). Our observation that riluzole reduced C3-expressing astrocytes while increasing S100A10-positive astrocytes indicates a shift toward a neuroprotective astrocyte profile rather than suppression of astrocytic activation.

Astrocyte morphology further reflects functional state. Simplified astrocytic arbors are associated with elevated C3 expression and increased amyloid burden (Liddelow et al., 2017; Chandra et al., 2023), whereas highly branched astrocytes establish extensive plaque contacts and are linked to reduced amyloid accumulation (Kraft et al., 2013). The restoration of astrocytic branching complexity observed with riluzole treatment therefore likely provides a structural substrate that supports enhanced Aβ handling and local synaptic protection.

Similarly, riluzole preserved microglial recruitment while biasing microglial responses toward a phagocytosis-competent, neuroprotective state. Although microglial activation has been viewed as detrimental in AD, accumulating evidence indicates that appropriate microglial recruitment can constrain amyloid plaque growth. Experimental depletion of microglia increases plaque size, whereas activation of TREM2 enhances microglial clustering around plaques and reduces amyloid burden (Zhao et al., 2017; Price et al., 2020).

The functional outcome of microglial activation depends on the balance between inflammatory and anti-inflammatory programs. Proinflammatory cytokine production is inversely correlated with the capacity of microglia to clear Aβ and can indirectly enhance amyloid deposition (Hickman et al., 2008; Pan et al., 2011; Heneka et al., 2013; Cheng et al., 2020). In contrast, microglia biased toward an anti-inflammatory, M2-like phenotype exhibit enhanced Aβ uptake and degradation (Heneka et al., 2013; Cherry et al., 2015; Fu et al., 2016). Consistent with this framework, riluzole reduced proinflammatory markers, promoted M2-like features, and normalized CD68 reactivity, suggesting restoration of efficient phagocytic function rather than suppression of microglial activity.

Morphological changes further reflected this functional shift. Aβ oligomer exposure increased dystrophic, hypofunctional microglia and reduced ramified surveillance states (Martini et al., 2020). Riluzole reversed this pattern, decreasing dystrophic microglia while increasing hypertrophic, injury-responsive microglia with enhanced branching complexity. Because increased microglial structural complexity correlates with improved Aβ uptake and clearance (Dodiya et al., 2019; Reichenbach et al., 2019), these morphological changes likely support effective amyloid handling.

## Conclusion

Taken together, our findings indicate that the beneficial effects of riluzole in Aβ oligomer pathology arise from coordinated modulation of neuron–glia interactions rather than from a single cellular target. By stabilizing glutamatergic signaling, riluzole simultaneously reduced neuronal vulnerability, preserved synaptic structure, and reshaped astrocytic and microglial responses toward states compatible with efficient amyloid handling and circuit maintenance. Importantly, riluzole maintained glial recruitment while restoring functional balance and morphological complexity, enabling effective Aβ clearance without exacerbating complement-mediated synaptic loss.

This convergence of neuronal protection, synaptic preservation, and adaptive glial responses provides a mechanistic framework linking glutamatergic modulation to circuit-level and behavioral resilience in the context of soluble Aβ toxicity. Our findings highlight glial functional state as a critical determinant of synaptic and cognitive outcomes and support the therapeutic potential of targeting neuron–glia interactions to restore network integrity in AD.

## Acknowledgments

This project was supported by Grants-in-Aid for Scientific Research C (D.M. Grant No. 23K06348), Scientific Research B (D.M. Grants No. 16H05129 and 19H03402), and Scientific Research in Innovative Areas (D.M. Grant No. 26115518), from the Ministry of Education, Culture, Sports, Science, and Technology of Japan. This project was also supported by the YU AI project of the Center for Information and Data Science Education, and JST SPRING, Grant Number JPMJSP2111.

## Notes

**Conflict of interest statement:** None of the authors has any conflict of interest to disclose. We confirm that we have read the Journal’s position on issues involved in ethical publication, and we affirm that this report is consistent with those guidelines. The funders had no role in study design, data collection and analysis, decision to publish, or preparation of the manuscript.

### Competing Interest Statement

The authors have declared no competing interest.

### Summary of Updates

We revised Figures 4F and 4G to strengthen the link between our findings and individual learning performance.

## References

Almeida, C. G., Tampellini, D., Takahashi, R. H., Greengard, P., Lin, M. T., Snyder, E. M., & Gouras, G. K. (2005). Beta-amyloid accumulation in APP mutant neurons reduces PSD-95 and GluR1 in synapses. Neurobiology of Disease, 20(2), 187–198. 10.1016/j.nbd.2005.02.008.

Bordji, K., Becerril-Ortega, J., Nicole, O., & Buisson, A. (2010). Activation of extrasynaptic, but not synaptic, NMDA receptors modifies amyloid precursor protein expression pattern and increases amyloid-β production. The Journal of Neuroscience, 30(47), 15927–15942. 10.1523/JNEUROSCI.3021-10.2010.

Brouillette, J., Caillierez, R., Zommer, N., Alves-Pires, C., Benilova, I., Blum, D., De Strooper, B., & Buee, L. (2012). Neurotoxicity and memory deficits induced by soluble low-molecular-weight amyloid- 1-42 oligomers are revealed in vivo by using a novel animal model. Journal of Neuroscience, 32(23), 7852–7861. 10.1523/JNEUROSCI.5901-11.2012.

Calvo-Rodriguez, M., Hou, S. S., Snyder, A. C., Kharitonova, E. K., Russ, A. N., Das, S., Fan, Z., Muzikansky, A., Garcia-Alloza, M., Serrano-Pozo, A., Hudry, E., & Bacskai, B. J. (2020). Increased mitochondrial calcium levels associated with neuronal death in a mouse model of Alzheimer’s disease. Nature Communications, 11(1), 2146. 10.1038/s41467-020-16074-2.

Chandra, S., Di Meco, A., Dodiya, H. B., Popovic, J., Cuddy, L. K., Weigle, I. Q., Zhang, X., Sadleir, K., Sisodia, S., & Vassar, R. (2023). The gut microbiome regulates astrocyte reaction to Aβ amyloidosis through microglial dependent and independent mechanisms. Molecular Neurodegeneration, 18(1), 45. 10.1186/s13024-023-00635-2.

Cherry, J. D., Olschowka, J. A., & O’Banion, M. K. (2015). Arginase 1+ microglia reduce Aβ plaque deposition during IL-1β-dependent neuroinflammation. Journal of Neuroinflammation, 12(1), 203. 10.1186/s12974-015-0411-8.

Cruz-Sanchez, A., Dematagoda, S., Ahmed, R., Mohanathaas, S., Odenwald, N., & Arruda-Carvalho, M. (2020). Developmental onset distinguishes three types of spontaneous recognition memory in mice. Scientific Reports, 10(1), 10612. 10.1038/s41598-020-67619-w.

Dickerson, B. C., Salat, D. H., Greve, D. N., Chua, E. F., Rand-Giovannetti, E., Rentz, D. M., Bertram, L., Mullin, K., Tanzi, R. E., Blacker, D., Albert, M. S., & Sperling, R. A. (2005). Increased hippocampal activation in mild cognitive impairment compared to normal aging and AD. Neurology, 65(3), 404–411. 10.1212/01.wnl.0000171450.97464.49.

Dodiya, H. B., Kuntz, T., Shaik, S. M., Baufeld, C., Leibowitz, J., Zhang, X., Gottel, N., Zhang, X., Butovsky, O., Gilbert, J. A., & Sisodia, S. S. (2019). Sex-specific effects of microbiome perturbations on cerebral Aβ amyloidosis and microglia phenotypes. Journal of Experimental Medicine, 216(7), 1542–1560. 10.1084/jem.20182386.

El-Khatib, S. M., Vagadia, A. R., Le, A. C. D., Baulch, J. E., Ng, D. Q., Du, M., Johnston, K. G., Tan, Z., Xu, X., Chan, A., & Acharya, M. (2024). BDNF augmentation reverses cranial radiation therapy-induced cognitive decline and neurodegenerative consequences. Acta Neuropathologica Communications, 12(1), 190. 10.1186/s40478-024-01906-9.

Feng, S., Wu, C., Zou, P., Deng, Q., Chen, Z., Li, M., Zhu, L., Li, F., Liu, T. C.-Y., Duan, R., & Yang, L. (2023). High-intensity interval training ameliorates Alzheimer’s disease-like pathology by regulating astrocyte phenotype-associated AQP4 polarization. Theranostics, 13(10), 3434–3450. 10.7150/thno.81951.

Forny-Germano, L., Lyra E Silva, N. M., Batista, A. F., Brito-Moreira, J., Gralle, M., Boehnke, S. E., Coe, B. C., Lablans, A., Marques, S. A., Martinez, A. M. B., Klein, W. L., Houzel, J.-C., Ferreira, S. T., Munoz, D. P., & De Felice, F. G. (2014). Alzheimer’s disease-like pathology induced by amyloid-β oligomers in nonhuman primates. The Journal of Neuroscience, 34(41), 13629–13643. 10.1523/JNEUROSCI.1353-14.2014.

Fu, A. K. Y., Hung, K.-W., Yuen, M. Y. F., Zhou, X., Mak, D. S. Y., Chan, I. C. W., Cheung, T. H., Zhang, B., Fu, W.-Y., Liew, F. Y., & Ip, N. Y. (2016). IL-33 ameliorates Alzheimer’s disease-like pathology and cognitive decline. Proceedings of the National Academy of Sciences, 113(19). 10.1073/pnas.1604032113.

Fumagalli, E., Funicello, M., Rauen, T., Gobbi, M., & Mennini, T. (2008). Riluzole enhances the activity of glutamate transporters GLAST, GLT1 and EAAC1. European Journal of Pharmacology, 578(2–3), 171–176. 10.1016/j.ejphar.2007.10.023.

Hardy, J., & Selkoe, D. J. (2002). The Amyloid hypothesis of Alzheimer’s disease: Progress and problems on the road to therapeutics. Science, 297(5580), 353–356. 10.1126/science.1072994.

Hascup, K. N., Findley, C. A., Britz, J., Esperant-Hilaire, N., Broderick, S. O., Delfino, K., Tischkau, S., Bartke, A., & Hascup, E. R. (2021). Riluzole attenuates glutamatergic tone and cognitive decline in AβPP/PS1 mice. Journal of Neurochemistry, 156(4), 513–523. 10.1111/jnc.15224.

Hascup, K. N., & Hascup, E. R. (2016). Soluble amyloid-β42 stimulates glutamate release through activation of the α7 nicotinic acetylcholine receptor. Journal of Alzheimer’s Disease, 53(1), 337–347. 10.3233/JAD-160041

Hector, A., & Brouillette, J. (2021). Hyperactivity induced by soluble amyloid-β oligomers in the early stages of Alzheimer’s disease. Frontiers in Molecular Neuroscience, 13, 600084. 10.3389/fnmol.2020.600084.

Heneka, M. T., Kummer, M. P., Stutz, A., Delekate, A., Schwartz, S., Vieira-Saecker, A., Griep, A., Axt, D., Remus, A., Tzeng, T.-C., Gelpi, E., Halle, A., Korte, M., Latz, E., & Golenbock, D. T. (2013). NLRP3 is activated in Alzheimer’s disease and contributes to pathology in APP/PS1 mice. Nature, 493(7434), 674–678. 10.1038/nature11729.

Hickman, S. E., Allison, E. K., & El Khoury, J. (2008). Microglial dysfunction and defective β-amyloid clearance pathways in aging Alzheimer’s disease mice. The Journal of Neuroscience, 28(33), 8354–8360. 10.1523/JNEUROSCI.0616-08.2008.

Hong, S., Beja-Glasser, V. F., Nfonoyim, B. M., Frouin, A., Li, S., Ramakrishnan, S., Merry, K. M., Shi, Q., Rosenthal, A., Barres, B. A., Lemere, C. A., Selkoe, D. J., & Stevens, B. (2016). Complement and microglia mediate early synapse loss in Alzheimer mouse models. Science, 352(6286), 712–716. 10.1126/science.aad8373.

Hunsberger, H. C., Weitzner, D. S., Rudy, C. C., Hickman, J. E., Libell, E. M., Speer, R., Gerhardt, G. A., & Reed, M. N. (2015). Riluzole rescues glutamate alterations, cognitive deficits, and tau pathology associated with P301L tau expression. Journal of Neurochemistry, 135(2), 381–394. 10.1111/jnc.13230.

Kim, J. Y., Mo, H., Kim, J., Kim, J. W., Nam, Y., Rim, Y. A., & Ju, J. H. (2022). Mitigating effect of estrogen in Alzheimer’s disease-mimicking cerebral organoid. Frontiers in Neuroscience, 16, 816174. 10.3389/fnins.2022.816174.

Knobloch, M., & Mansuy, I. M. (2008). Dendritic spine loss and synaptic alterations in Alzheimer’s disease. Molecular Neurobiology, 37(1), 73–82. 10.1007/s12035-008-8018-z.

Koh, M. T., Haberman, R. P., Foti, S., McCown, T. J., & Gallagher, M. (2010). Treatment strategies targeting excess hippocampal activity benefit aged rats with cognitive impairment. Neuropsychopharmacology, 35(4), 1016–1025. 10.1038/npp.2009.207.

Kraft, A. W., Hu, X., Yoon, H., Yan, P., Xiao, Q., Wang, Y., Gil, S. C., Brown, J., Wilhelmsson, U., Restivo, J. L., Cirrito, J. R., Holtzman, D. M., Kim, J., Pekny, M., & Lee, J. (2013). Attenuating astrocyte activation accelerates plaque pathogenesis in APP/PS1 mice. The FASEB Journal, 27(1), 187–198. 10.1096/fj.12-208660.

Kyllo, T., Allocco, D., Hei, L. V., Wulff, H., & Erickson, J. D. (2024). Riluzole attenuates acute neural injury and reactive gliosis, hippocampal-dependent cognitive impairments and spontaneous recurrent generalized seizures in a rat model of temporal lobe epilepsy. Frontiers in Pharmacology, 15, 1466953. 10.3389/fphar.2024.1466953.

Lacor, P. N., Buniel, M. C., Chang, L., Fernandez, S. J., Gong, Y., Viola, K. L., Lambert, M. P., Velasco, P. T., Bigio, E. H., Finch, C. E., Krafft, G. A., & Klein, W. L. (2004). Synaptic targeting by Alzheimer’s-related amyloid β oligomers. The Journal of Neuroscience, 24(45), 10191–10200. 10.1523/JNEUROSCI.3432-04.2004.

Lacor, P. N., Buniel, M. C., Furlow, P. W., Sanz Clemente, A., Velasco, P. T., Wood, M., Viola, K. L., & Klein, W. L. (2007). Aβ oligomer-induced aberrations in synapse composition, shape, and density provide a molecular basis for loss of connectivity in Alzheimer’s disease. The Journal of Neuroscience, 27(4), 796–807. 10.1523/JNEUROSCI.3501-06.2007.

Lamanauskas, N., & Nistri, A. (2008). Riluzole blocks persistent Na^+^ and Ca^2+^ currents and modulates release of glutamate via presynaptic NMDA receptors on neonatal rat hypoglossal motoneurons *in vitro*. European Journal of Neuroscience, 27(10), 2501–2514. 10.1111/j.1460-9568.2008.06211.x.

Lee, C. Y. D., & Landreth, G. E. (2010). The role of microglia in amyloid clearance from the AD brain. Journal of Neural Transmission, 117(8), 949–960. 10.1007/s00702-010-0433-4.

Lei, M., Xu, H., Li, Z., Wang, Z., O’Malley, T. T., Zhang, D., Walsh, D. M., Xu, P., Selkoe, D. J., & Li, S. (2016). Soluble Aβ oligomers impair hippocampal LTP by disrupting glutamatergic/GABAergic balance. Neurobiology of Disease, 85, 111–121. 10.1016/j.nbd.2015.10.019.

Li, S., Jin, M., Koeglsperger, T., Shepardson, N. E., Shankar, G. M., & Selkoe, D. J. (2011). Soluble Aβ oligomers inhibit long-term potentiation through a mechanism involving excessive activation of extrasynaptic NR2B-containing NMDA receptors. The Journal of Neuroscience, 31(18), 6627–6638. 10.1523/JNEUROSCI.0203-11.2011.

Lian, H., Litvinchuk, A., Chiang, A. C.-A., Aithmitti, N., Jankowsky, J. L., & Zheng, H. (2016). Astrocyte-microglia cross talk through complement activation modulates amyloid pathology in mouse models of Alzheimer’s disease. The Journal of Neuroscience, 36(2), 577–589. 10.1523/JNEUROSCI.2117-15.2016.

Liddelow, S. A., Guttenplan, K. A., Clarke, L. E., Bennett, F. C., Bohlen, C. J., Schirmer, L., Bennett, M. L., Münch, A. E., Chung, W.-S., Peterson, T. C., Wilton, D. K., Frouin, A., Napier, B. A., Panicker, N., Kumar, M., Buckwalter, M. S., Rowitch, D. H., Dawson, V. L., Dawson, T. M., … Barres, B. A. (2017). Neurotoxic reactive astrocytes are induced by activated microglia. Nature, 541(7638), 481–487. 10.1038/nature21029.

Lv, Z., Chen, L., Chen, P., Peng, H., Rong, Y., Hong, W., Zhou, Q., Li, N., Li, B., Paolicelli, R. C., & Zhan, Y. (2024). Clearance of β-amyloid and synapses by the optogenetic depolarization of microglia is complement selective. Neuron, 112(5), 740–754.e7. 10.1016/j.neuron.2023.12.003.

Martini, A. C., Helman, A. M., McCarty, K. L., Lott, I. T., Doran, E., Schmitt, F. A., & Head, E. (2020). Distribution of microglial phenotypes as a function of age and Alzheimer’s disease neuropathology in the brains of people with Down syndrome. *Alzheimer’s & Dementia: Diagnosis*, Assessment & Disease Monitoring, 12(1). 10.1002/dad2.12113.

Mehder, R. H., Bennett, B. M., & Andrew, R. D. (2020). Morphometric analysis of hippocampal and neocortical pyramidal neurons in a mouse model of late onset Alzheimer’s disease. Journal of Alzheimer’s Disease, 74(4), 1069–1083. 10.3233/JAD-191067.

Min-Kaung-Wint-Mon, Kida, H., Kanehisa, I., Kurose, M., Ishikawa, J., Sakimoto, Y., Paw-Min-Thein-Oo, Kimura, R., & Mitsushima, D. (2024). Adverse effects of Aβ1-42 Oligomers: Impaired contextual memory and altered intrinsic properties of CA1 pyramidal neurons. Biomolecules, 14(11), 1425. 10.3390/biom14111425.

Mitsushima, D., Ishihara, K., Sano, A., Kessels, H. W., & Takahashi, T. (2011). Contextual learning requires synaptic AMPA receptor delivery in the hippocampus. Proceedings of the National Academy of Sciences, 108(30), 12503–12508. 10.1073/pnas.1104558108.

Mizuta, I., Ohta, M., Ohta, K., Nishimura, M., Mizuta, E., & Kuno, S. (2001). Riluzole stimulates nerve growth factor, brain-derived neurotrophic factor and glial cell line-derived neurotrophic factor synthesis in cultured mouse astrocytes. Neuroscience Letters, 310(2–3), 117–120. 10.1016/S0304-3940(01)02098-5.

Okamoto, M., Gray, J. D., Larson, C. S., Kazim, S. F., Soya, H., McEwen, B. S., & Pereira, A. C. (2018). Riluzole reduces amyloid beta pathology, improves memory, and restores gene expression changes in a transgenic mouse model of early-onset Alzheimer’s disease. Translational Psychiatry, 8(1), 153. 10.1038/s41398-018-0201-z.

Pan, X., Zhu, Y., Lin, N., Zhang, J., Ye, Q., Huang, H., & Chen, X. (2011). Microglial phagocytosis induced by fibrillar β-amyloid is attenuated by oligomeric β-amyloid: Implications for Alzheimer’s disease. Molecular Neurodegeneration, 6(1), 45. 10.1186/1750-1326-6-45.

Patnaik, A., Zagrebelsky, M., Korte, M., & Holz, A. (2020). Signaling via the p75 neurotrophin receptor facilitates amyloid β-induced dendritic spine pathology. Scientific Reports, 10(1), 13322. 10.1038/s41598-020-70153-4.

Pereira, A. C., Gray, J. D., Kogan, J. F., Davidson, R. L., Rubin, T. G., Okamoto, M., Morrison, J. H., & McEwen, B. S. (2017). Age and Alzheimer’s disease gene expression profiles reversed by the glutamate modulator riluzole. Molecular Psychiatry, 22(2), 296–305. 10.1038/mp.2016.33.

Pereira, A. C., Lambert, H. K., Grossman, Y. S., Dumitriu, D., Waldman, R., Jannetty, S. K., Calakos, K., Janssen, W. G., McEwen, B. S., & Morrison, J. H. (2014). Glutamatergic regulation prevents hippocampal-dependent age-related cognitive decline through dendritic spine clustering. Proceedings of the National Academy of Sciences, 111(52), 18733–18738. 10.1073/pnas.1421285111.

Price, B. R., Sudduth, T. L., Weekman, E. M., Johnson, S., Hawthorne, D., Woolums, A., & Wilcock, D. M. (2020). Therapeutic TREM2 activation ameliorates amyloid-beta deposition and improves cognition in the 5XFAD model of amyloid deposition. Journal of Neuroinflammation, 17(1), 238. 10.1186/s12974-020-01915-0.

Reichenbach, N., Delekate, A., Plescher, M., Schmitt, F., Krauss, S., Blank, N., Halle, A., & Petzold, G. C. (2019). Inhibition of Stat3-mediated astrogliosis ameliorates pathology in an Alzheimer’s disease model. EMBO Molecular Medicine, 11(2), e9665. 10.15252/emmm.201809665.

Ren, S., Chen, P., Jiang, H., Mi, Z., Xu, F., Hu, B., Zhang, J., & Zhu, Z. (2014). Persistent sodium currents contribute to Aβ_1-42_-induced hyperexcitation of hippocampal CA1 pyramidal neurons. Neuroscience Letters, 580, 62–67. 10.1016/j.neulet.2014.07.050.

Rostami, J., Mothes, T., Kolahdouzan, M., Eriksson, O., Moslem, M., Bergström, J., Ingelsson, M., O’Callaghan, P., Healy, L. M., Falk, A., & Erlandsson, A. (2021). Crosstalk between astrocytes and microglia results in increased degradation of α-synuclein and amyloid-β aggregates. Journal of Neuroinflammation, 18(1), 124. 10.1186/s12974-021-02158-3.

Sakimoto, Y., Shintani, A., Yoshiura, D., Goshima, M., Kida, H., & Mitsushima, D. (2022). A critical period for learning and plastic changes at hippocampal CA1 synapses. Scientific Reports, 12(1), 7199. 10.1038/s41598-022-10453-z.

Serrano-Pozo, A., Muzikansky, A., Gómez-Isla, T., Growdon, J. H., Betensky, R. A., Frosch, M. P., & Hyman, B. T. (2013). Differential relationships of reactive astrocytes and microglia to fibrillar amyloid deposits in Alzheimer disease. Journal of Neuropathology & Experimental Neurology, 72(6), 462–471. 10.1097/NEN.0b013e3182933788.

Shen, Y., Zhang, X., Liu, S., Xin, L., Xuan, W., Zhuang, C., Chen, Y., Chen, B., Zheng, X., Wu, R., & Lin, Y. (2025). CEST imaging combined with 1H-MRS reveal the neuroprotective effects of riluzole by improving neurotransmitter imbalances in Alzheimer’s disease mice. Alzheimer’s Research & Therapy, 17(1), 20. 10.1186/s13195-025-01672-3.

Šišková, Z., Justus, D., Kaneko, H., Friedrichs, D., Henneberg, N., Beutel, T., Pitsch, J., Schoch, S., Becker, A., von der Kammer, H., & Remy, S. (2014). Dendritic structural degeneration is functionally linked to cellular hyperexcitability in a mouse model of Alzheimer’s disease. Neuron, 84(5), 1023–1033. 10.1016/j.neuron.2014.10.024.

Stine, W. B., Dahlgren, K. N., Krafft, G. A., & LaDu, M. J. (2003). In vitro characterization of conditions for amyloid-β peptide oligomerization and fibrillogenesis. Journal of Biological Chemistry, 278(13), 11612–11622. 10.1074/jbc.M210207200.

Tu, S., Okamoto, S., Lipton, S. A., & Xu, H. (2014). Oligomeric Aβ-induced synaptic dysfunction in Alzheimer’s disease. Molecular Neurodegeneration, 9(1), 48. 10.1186/1750-1326-9-48.

Wei, W., Nguyen, L. N., Kessels, H. W., Hagiwara, H., Sisodia, S., & Malinow, R. (2010). Amyloid beta from axons and dendrites reduces local spine number and plasticity. Nature Neuroscience, 13(2), 190–196. 10.1038/nn.2476

Wu, Y., Satkunendrarajah, K., Teng, Y., Chow, D. S.-L., Buttigieg, J., & Fehlings, M. G. (2013). Delayed post-injury administration of riluzole is neuroprotective in a preclinical rodent model of cervical spinal cord injury. Journal of Neurotrauma, 30(6), 441–452. 10.1089/neu.2012.2622.

Yang, Y., Ji, W., Zhang, Y., Zhou, L., Chen, H., Yang, N., & Zhu, Z. (2021). Riluzole ameliorates soluble Aβ1–42-induced impairments in spatial memory by modulating the glutamatergic/GABAergic balance in the dentate gyrus. Progress in Neuro-Psychopharmacology and Biological Psychiatry, 108, 110077. 10.1016/j.pnpbp.2020.110077.

Zarate, C. A., & Manji, H. K. (2008). Riluzole in psychiatry: A systematic review of the literature. Expert Opinion on Drug Metabolism & Toxicology, 4(9), 1223–1234. 10.1517/17425255.4.9.1223.

Zhao, R., Hu, W., Tsai, J., Li, W., & Gan, W.-B. (2017). Microglia limit the expansion of β-amyloid plaques in a mouse model of Alzheimer’s disease. Molecular Neurodegeneration, 12(1), 47. 10.1186/s13024-017-0188-6.

Zhong, F., Liu, L., Wei, J.-L., & Dai, R.-P. (2019). Step by step Golgi-Cox staining for cryosection. Frontiers in Neuroanatomy, 13, 62. 10.3389/fnana.2019.00062.

Zott, B., Simon, M. M., Hong, W., Unger, F., Chen-Engerer, H.-J., Frosch, M. P., Sakmann, B., Walsh, D. M., & Konnerth, A. (2019). A vicious cycle of β amyloid–dependent neuronal hyperactivation. Science, 365(6453), 559–565. 10.1126/science.aay0198.

