## Supplementary Information for "Riluzole shifts glial responses to protect synapses and memory in Aβ oligomer–treated rats"

**Table S1** Antibodies used in this study. The table summarizes information about host species, concentration, supplier company, and identifier.

| Antibody | Dilution | Supplier | Identifier |
|---|---|---|---|
| Mouse anti-NeuN | 1:250 | Invitrogen | 18-7373 |
| Mouse anti-MAP2 | 1:500 | Sigma-Aldrich | M9942 |
| Mouse anti-C3 | 1:100 | Santa Cruz Biotechnology | SC-28294 |
| Mouse anti-S100A10 | 1:100 | Cell Signaling Technology | 5529T |
| Mouse anti-CD11b | 1:200 | Sigma-Aldrich | CBL1512-25UG |
| Mouse anti-CD68 | 1:200 | Abcam | ab955 |
| Mouse anti-iNos2 | 1:50 | Santa Cruz Biotechnology | SC-7271 |
| Mouse anti-arginase1 | 1:100 | Santa Cruz Biotechnology | SC-271430 |
| Mouse anti-PSD95 | 1:500 | Neuromab | 750-28 |
| Rabbit anti-cleaved caspase 3 | 1:100 | Cell Signaling Technology | 9664T |
| Rabbit anti-PSD95 | 1:500 | Abcam | ab18258 |
| Rabbit anti-GFAP | 1:1000 | Dako | Z0334 |
| Rabbit anti-Iba1 | 1:250 | Wako | 019-19741 |
| Guinea pig anti-vGlut1 | 1:1000 | Wako | 013-2811 |
| Lecanemab | 1:500 | Selleck | A3112 |
| Goat anti-mouse Alexa 594 | 1:500 | Abcam | ab150116 |
| Goat anti-mouse Alexa FITC | 1:500 | Jackson ImmunoResearch | F-2761 |
| Goat anti-rabbit Texas Red | 1:500 | Jackson ImmunoResearch | 72626 |
| Goat anti-rabbit FITC | 1:500 | Chemicon | AP307F |
| Goat anti- guinea pig Alexa 488 | 1:500 | Abcam | ab150185 |
| Goat anti-human Alexa 488 | 1:500 | Invitrogen | A56869 |

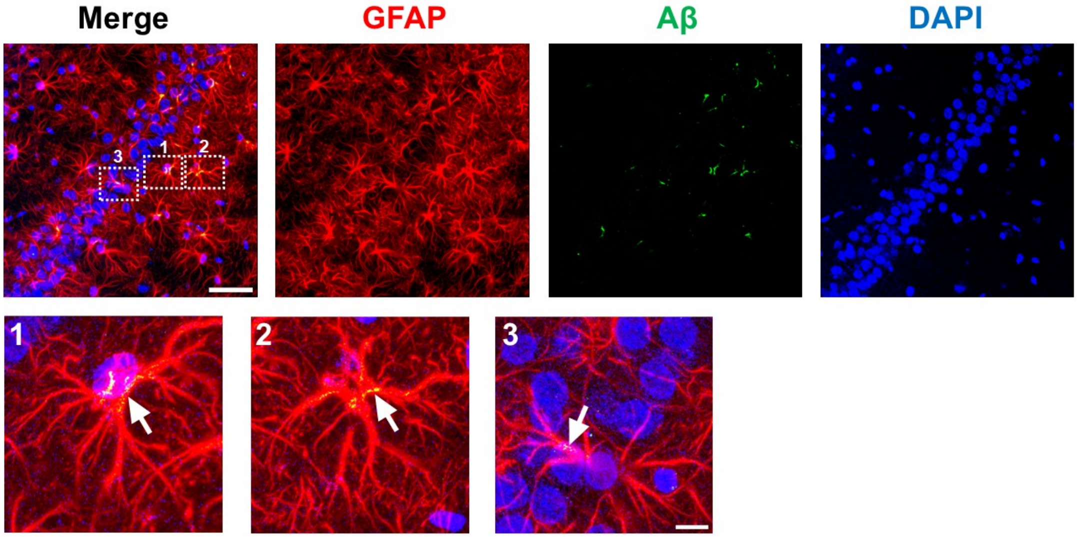


**Figure S1. Effect of riluzole on the engulfment of $A\beta_{1-42}$ oligomers by astrocytes in the dorsal CA1 region.** Representative confocal z-stack (4 μm) images (upper) and zoomed-in images (lower) showing the engulfment of $A\beta_{1-42}$ oligomers by astrocytes in the dorsal CA1 region. The images were 3D rendered in MIP mode using Imaris software (v.10.0.2; Bitplane). The magnification is ×400, and the arrows indicate engulfed Aβ. Scale bars: 50 μm (upper) and 10 μm (lower).

**Table S2** Summary of statistical results from **Figure 5**

| Parameter | ANOVA | Bonferroni's test |
|---|---|---|
| **Figure 5D** | | |
| Total spines (per 20 μm) | Aβ: $F_{1,67} = 75.174$, $p < 0.0001$<br>Riluzole: $F_{1,67} = 67.004$, $p < 0.0001$<br>Interaction: $F_{1,67} = 75.174$, $p < 0.0001$ | Aβ *vs*. saline, $p < 0.0001$<br>Aβ *vs*. Aβ+RLZ, $p < 0.0001$ |
| Thin spines | Aβ: $F_{1,67} = 1.448$, $p = 0.2331$<br>Riluzole: $F_{1,67} = 6.564$, $p = 0.0127$<br>Interaction: $F_{1,67} = 4.313$, $p = 0.0417$ | Aβ *vs*. saline, $p = 0.1253$<br>Aβ *vs*. Aβ+RLZ, $p = 0.0081$ |
| Mushroom spines | Aβ: $F_{1,67} = 38.418$, $p < 0.0001$<br>Riluzole: $F_{1,67} = 23.657$, $p < 0.0001$<br>Interaction: $F_{1,67} = 17.612$, $p < 0.0001$ | Aβ *vs*. saline, $p < 0.0001$<br>Aβ *vs*. Aβ+RLZ, $p < 0.0001$ |
| Stubby spines | Aβ: $F_{1,67} =37.536$, $p < 0.0001$<br>Riluzole: $F_{1,67} = 21.515$, $p < 0.0001$<br>Interaction: $F_{1,67} = 30.322$, $p < 0.0001$ | Aβ *vs*. saline, $p < 0.0001$<br>Aβ *vs*. Aβ+RLZ, $p < 0.0001$ |
| **Figure 5E** | | |
| Head area ($\mu m^2$) | Aβ: $F_{1,428} = 8.600$, $p = 0.0035$<br>Riluzole: $F_{1,428} = 2.820$, $p = 0.0938$<br>Interaction: $F_{1,428} = 5.786$, $p = 0.0166$ | Aβ *vs*. saline, $p = 0.0010$<br>Aβ *vs*. Aβ+RLZ, $p = 0.0224$ |
| Neck length (μm) | Aβ: $F_{1,428} = 1.898$, $p = 0.1690$<br>Riluzole: $F_{1,428} = 7.642$, $p = 0.0059$<br>Interaction: $F_{1,430} = 8.125$, $p = 0.0046$ | Aβ *vs*. saline, $p = 0.0165$<br>Aβ *vs*. Aβ+RLZ, $p = 0.0004$ |

| | | |
|---|---|---|
| Neck width (µm) | Aβ: $F_{1, 428}$ =, $p < 0.0001$<br>Riluzole: $F_{1, 428} = 67.004$, $p < 0.0001$<br>Interaction: $F_{1, 428} = 75.174$, $p < 0.0001$ | Aβ *vs*. saline, $p = 0.7588$<br>Aβ *vs*. Aβ+RLZ, $p = 0.2832$ |
| **Figure 5F** | | |
| Total spines (per 20 µm) | Aβ: $F_{1, 79} = 114.832$, $p < 0.0001$<br>Riluzole: $F_{1, 79} = 67.322$, $p < 0.0001$<br>Interaction: $F_{1, 79} = 99.202$, $p < 0.0001$ | Aβ *vs*. saline, $p < 0.0001$<br>Aβ *vs*. Aβ+RLZ, $p < 0.0001$ |
| Thin spines | Aβ: $F_{1, 79} = 33.145$, $p < 0.0001$<br>Riluzole: $F_{1, 79} = 6.343$, $p = 0.0138$<br>Interaction: $F_{1, 79} = 17.594$, $p < 0.0001$ | Aβ *vs*. saline, $p < 0.0001$<br>Aβ *vs*. Aβ+RLZ, $p < 0.0001$ |
| Mushroom spines | Aβ: $F_{1, 79} = 7.307$, $p = 0.0084$<br>Riluzole: $F_{1, 79} = 15.504$, $p = 0.0002$<br>Interaction: $F_{1, 79} = 7.449$, $p = 0.0078$ | Aβ *vs*. saline, $p = 0.0014$<br>Aβ *vs*. Aβ+RLZ, $p < 0.0001$ |
| Stubby spines | Aβ: $F_{1, 79} = 19.595$, $p < 0.0001$<br>Riluzole: $F_{1, 79} = 10.161$, $p = 0.0021$<br>Interaction: $F_{1, 79} = 18.412$, $p < 0.0001$ | Aβ *vs*. saline, $p < 0.0001$<br>Aβ *vs*. Aβ+RLZ, $p < 0.0001$ |
| **Figure 5G** | | |
| Head area ($\mu m^2$) | Aβ: $F_{1, 422} = 40.479$, $p < 0.0001$<br>Riluzole: $F_{1, 422} = 14.006$, $p = 0.0002$<br>Interaction: $F_{1, 422} = 25.900$, $p < 0.0001$ | Aβ *vs*. saline, $p < 0.0001$<br>Aβ *vs*. Aβ+RLZ, $p < 0.0001$ |
| Neck length (µm) | Aβ: $F_{1, 422} = 6.772$, $p = 0.0096$<br>Riluzole: $F_{1, 422} = 2.837$, $p = 0.0928$<br>Interaction: $F_{1, 422} = 4.646$, $p = 0.0317$ | Aβ *vs*. saline, $p = 0.0046$<br>Aβ *vs*. Aβ+RLZ, $p = 0.0438$ |
| Neck width (µm) | Aβ: $F_{1, 422}$ =45.320, $p < 0.0001$<br>Riluzole: $F_{1, 422} = 12.037$, $p = 0.0006$<br>Interaction: $F_{1, 422} = 25.244$, $p < 0.0001$ | Aβ *vs*. saline, $p < 0.0001$<br>Aβ *vs*. Aβ+RLZ, $p < 0.0001$ |
| **Figure 5H** | | |
| Total spines (per 20 µm) | Aβ: $F_{1, 75} = 26.326$, $p < 0.0001$<br>Riluzole: $F_{1, 75} = 25.505$, $p < 0.0001$<br>Interaction: $F_{1, 75} = 28.008$, $p < 0.0001$ | Aβ *vs*. saline, $p < 0.0001$<br>Aβ *vs*. Aβ+RLZ, $p < 0.0001$ |
| Thin spines | Aβ: $F_{1, 75} = 10.264$, $p = 0.0020$<br>Riluzole: $F_{1, 75} = 8.689$, $p = 0.0043$<br>Interaction: $F_{1, 75} = 10.570$, $p = 0.0017$ | Aβ *vs*. saline, $p < 0.0001$<br>Aβ *vs*. Aβ+RLZ, $p = 0.0002$ |
| Mushroom spines | Aβ: $F_{1, 75} = 9.188$, $p = 0.0033$<br>Riluzole: $F_{1, 75} = 3.254$, $p = 0.0753$<br>Interaction: $F_{1, 75} = 9.415$, $p = 0.0030$ | Aβ *vs*. saline, $p = 0.0002$<br>Aβ *vs*. Aβ+RLZ, $p = 0.0045$ |
| Stubby spines | Aβ: $F_{1, 75} = 0.017$, $p = 0.8962$<br>Riluzole: $F_{1, 75} = 3.044$, $p = 0.0852$<br>Interaction: $F_{1, 75} = 0.082$, $p = 0.776$ | Aβ *vs*. saline, $p > 0.9999$<br>Aβ *vs*. Aβ+RLZ, $p = 0.8828$ |
| **Figure 5I** | | |
| Head area ($\mu m^2$) | Aβ: $F_{1, 407} = 7.877$, $p = 0.0052$<br>Riluzole: $F_{1, 407} = 4.809$, $p = 0.0289$<br>Interaction: $F_{1, 407} = 2.564$, $p = 0.1101$ | Aβ *vs*. saline, $p = 0.0110$<br>Aβ *vs*. Aβ+RLZ, $p = 0.0434$ |
| Neck length (µm) | Aβ: $F_{1, 407} = 2.290$, $p = 0.1310$<br>Riluzole: $F_{1, 407} = 7.178$, $p = 0.0077$<br>Interaction: $F_{1, 407} = 9.573$, $p < 0.0021$ | Aβ *vs*. saline, $p = 0.0068$<br>Aβ *vs*. Aβ+RLZ, $p = 0.0003$ |
| Neck width (µm) | Aβ: $F_{1, 407}$ =18.177, $p < 0.0001$<br>Riluzole: $F_{1, 407} = 15.166$, $p = 0.0001$<br>Interaction: $F_{1, 407} = 8.049$, $p = 0.0048$ | Aβ *vs*. saline, $p < 0.0001$<br>Aβ *vs*. Aβ+RLZ, $p < 0.0001$ |

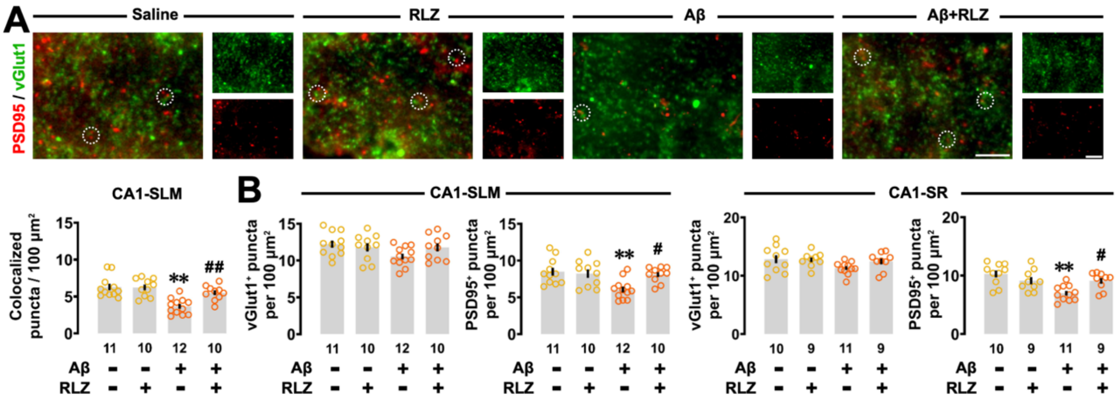

**Figure S2. Effects of riluzole on $A\beta_{1\text{-}42}$ oligomer-induced synaptic loss of the dorsal CA1 region. (A)** Representative fluorescence images and semi-quantitative analysis of colocalized vGlut1/PSD95 puncta per 100 μm$^2$ in the stratum lacunosum-moleculare of the dorsal region. The Aβ-induced reduction in synaptic puncta was reversed in the Aβ +RLZ group. **(B)** Quantitative analysis showed that Aβ oligomers did not significantly affect presynaptic vGlut1 puncta, but decreased total postsynaptic PSD95. This resulted in a decrease in colocalized synaptic puncta in the stratum radiatum and stratum lacunosum-moleculare of the dorsal CA1 region, respectively. Magnification is ×400 (digital zoom 3×), and the dotted circles indicate colocalized puncta. Scale bars: 5 μm. Data are plotted as individual points and expressed as mean ± SEM. ** $p < 0.01$ *vs.* saline. # $p < 0.05$, ## $p < 0.01$ *vs.* Aβ.

**Table S3** Summary of statistical results from **Figure S2**

| Parameter | ANOVA | Bonferroni's test |
|---|---|---|
| | **Figure S1A** | |
| Colocalized puncta per 100 μm$^2$ | Aβ: $F_{1,39} = 20.901, p < 0.0001$<br>Riluzole: $F_{1,39} = 6.032, p = 0.0186$<br>Interaction: $F_{1,39} = 6.573, p = 0.0143$ | Aβ *vs.* saline, $p < 0.0001$<br>Aβ *vs.* Aβ+RLZ, $p = 0.0055$ |
| | **Figure S1B** | |
| vGlut1$^+$ puncta (CA1-SR) | Aβ: $F_{1,35} = 3.270, p = 0.0791$<br>Riluzole: $F_{1,35} = 1.072, p = 0.3076$<br>Interaction: $F_{1,35} = 1.473, p = 0.2330$ | Aβ *vs.* saline, $p = 0.0196$<br>Aβ *vs.* Aβ+RLZ, $p = 0.7007$ |
| PSD95$^+$ puncta (CA1-SR) | Aβ: $F_{1,35} = 9.912, p = 0.0039$<br>Riluzole: $F_{1,35} = 0.8330, p = 0.3677$<br>Interaction: $F_{1,35} = 9.561, p = 0.0039$ | Aβ *vs.* saline, $p = 0.0003$<br>Aβ *vs.* Aβ+RLZ, $p = 0.0415$ |
| vGlut1$^+$ puncta (CA1-SLM) | Aβ: $F_{1,39} = 2.881, p = 0.0976$<br>Riluzole: $F_{1,39} = 0.664, p = 0.4200$<br>Interaction: $F_{1,39} = 2.835, p = 0.1002$ | Aβ *vs.* saline, $p = 0.1060$<br>Aβ *vs.* Aβ+RLZ, $p = 0.4921$ |
| PSD95$^+$ puncta (CA1-SLM) | Aβ: $F_{1,39} = 7.510, p = 0.0092$<br>Riluzole: $F_{1,39} = 3.548, p = 0.0671$<br>Interaction: $F_{1,39} = 6.022, p = 0.0187$ | Aβ *vs.* saline, $p = 0.0029$<br>Aβ *vs.* Aβ+RLZ, $p = 0.0216$ |

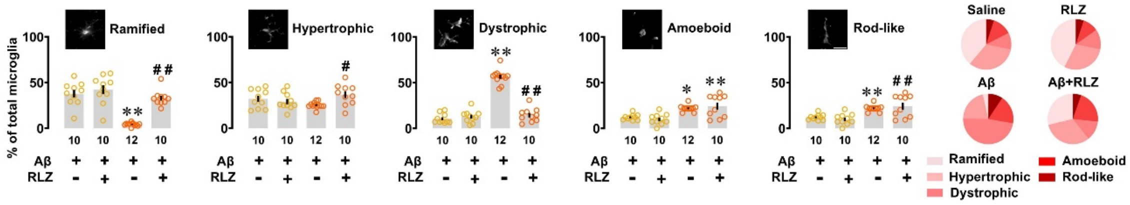


**Figure S3. Effect of riluzole on $A\beta_{1\text{-}42}$ oligomer-induced alteration in microglia phenotype in the dorsal CA1 region.** Analysis of the morphological classification of microglia showed that the reduction of ramified microglia and the increase in dystrophic and rod-like microglia percentage induced by Aβ were reversed in the Aβ + RLZ group. The amoeboid type remained unchanged. Magnification is ×400 (digital zoom 3×). Scale bar: 20 µm. The number of slices is shown at the bottom of each bar. Data are plotted as individual points and expressed as the mean ± SEM. * $p < 0.05$; ** $p < 0.01$ *vs.* saline. $^{\#}p < 0.05$, $^{\#\#}p < 0.01$ *vs.* Aβ.

**Table S4** Summary of statistical results from **Figure S3**

| Parameter | ANOVA | Bonferroni's test |
|---|---|---|
| Ramified | Aβ: $F_{1,38} = 41.357$, $p < 0.0001$<br>Riluzole: $F_{1,38} = 24.918$, $p < 0.0001$<br>Interaction: $F_{1,38} = 13.375$, $p = 0.0008$ | Aβ *vs.* saline, $p < 0.0001$<br>Aβ *vs.* Aβ+RLZ, $p < 0.0001$ |
| Hypertrophic | Aβ: $F_{1,38} = 0.011$, $p = 0.9158$<br>Riluzole: $F_{1,38} = 2.281$, $p = 0.1392$<br>Interaction: $F_{1,38} = 6.775$, $p = 0.0131$ | Aβ *vs.* saline, $p = 0.4741$<br>Aβ *vs.* Aβ+RLZ, $p = 0.0305$ |
| Dystrophic | Aβ: $F_{1,38} = 123.939$, $p < 0.0001$<br>Riluzole: $F_{1,38} = 85.776$, $p < 0.0001$<br>Interaction: $F_{1,38} =106.689$, $p < 0.0001$ | Aβ *vs.* saline, $p < 0.0001$<br>Aβ *vs.* Aβ+RLZ, $p < 0.0001$ |
| Amoeboid | Aβ: $F_{1,38} = 31.620$, $p < 0.0001$<br>Riluzole: $F_{1,38} = 0.0003$, $p = 0.9534$<br>Interaction: $F_{1,38} = 1.1565$, $p = 0.2890$ | Aβ *vs.* saline, $p = 0.0131$<br>saline *vs.* Aβ+RLZ, $p = 0.0021$ |
| Rod-like | Aβ: $F_{1,38} = 15.166$, $p = 0.0004$<br>Riluzole: $F_{1,38} = 6.273$, $p = 0.0167$<br>Interaction: $F_{1,38} = 7.726$, $p = 0.0084$ | Aβ *vs.* saline, $p = 0.0001$<br>Aβ *vs.* Aβ+RLZ, $p = 0.0029$ |
